# miRNA upregulate protein and glycan expression via direct activation in proliferating cells

**DOI:** 10.1101/2022.04.01.486772

**Authors:** Faezeh Jame-Chenarboo, Hoi Hei Ng, Dawn Macdonald, Lara K. Mahal

## Abstract

The dominant paradigm is that miRNA binding to mRNA represses protein expression. Activation by miRNA has been observed in select circumstances (quiescent cells, mitochondria), but is not thought a feature of miRNA action in actively dividing cells. Herein, we comprehensively map the miRNA regulation of α-2,6-sialyltransferases ST6GAL1 and ST6GAL2 using a high-throughput assay (miRFluR). We find the majority of miRNA targeting ST6GAL1, the main enzyme controlling α-2,6-sialylation, upregulate protein expression. In contrast, those that regulate ST6GAL2 are predominantly downregulatory. We provide evidence that miRNA-mediated upregulation occurs in proliferating cells and is a direct effect. Further, we show that AGO2 and FXR1 are required. Our data expands current understanding of miRNA, providing strong evidence of both upregulatory and downregulatory roles for these non-coding RNA.

**One-Sentence Summary:** miRNA directly activate expression of α-2,6-sialyltransferases and sialylation, expanding miRNA actions in dividing cells.

## Main Text

The canonical view of microRNAs (miRNAs, miRs) is that they are posttranscriptional repressors, binding the 3’-UTR of mRNA causing destabilization and/or loss of translation (*1*). In recent work Corey and coworkers hint at more complex regulation of protein expression by miRNAs, providing evidence that the miRNA machinery may be involved in promoting expression in dividing cells (*2*). Consistent with this, miRNAs can activate expression in select circumstances (e.g. senescent cells (*3*), oocytes (*4*) and mitochondria (*5*)) where the mRNA is destabilized, lacking a 5’-cap and a typical poly(A) tail. In actively dividing cells, mRNA do not meet these requirements and upregulation of expression by miRNA is not thought to occur.

Our laboratory works on regulation of glycosylation by miRNAs (*6-9*). Glycan regulation by miRNAs impacts multiple biological processes including melanoma metastasis (*10*), endothelial to hematopoetic transition (*11*), and epithelial-to-mesenchymal transition (*7*). α-2,6-sialic acid is one of the most studied glycosylation motifs, with clear roles in immunology, infectious disease, and cancer biology (*12, 13*). This modification is biosynthesized by two enzymes: ST6-beta-galactoside-α-2,6-sialyltranferase-1 (ST6GAL1), expressed throughout the human body, and ST6-beta-galactoside-α-2,6-sialyltranferase-2 (ST6GAL2), predominantly seen in breast and brain (**Fig. 1A**) (*14, 15*). Little is known about their regulation by miRNA.

**Fig. 1.**
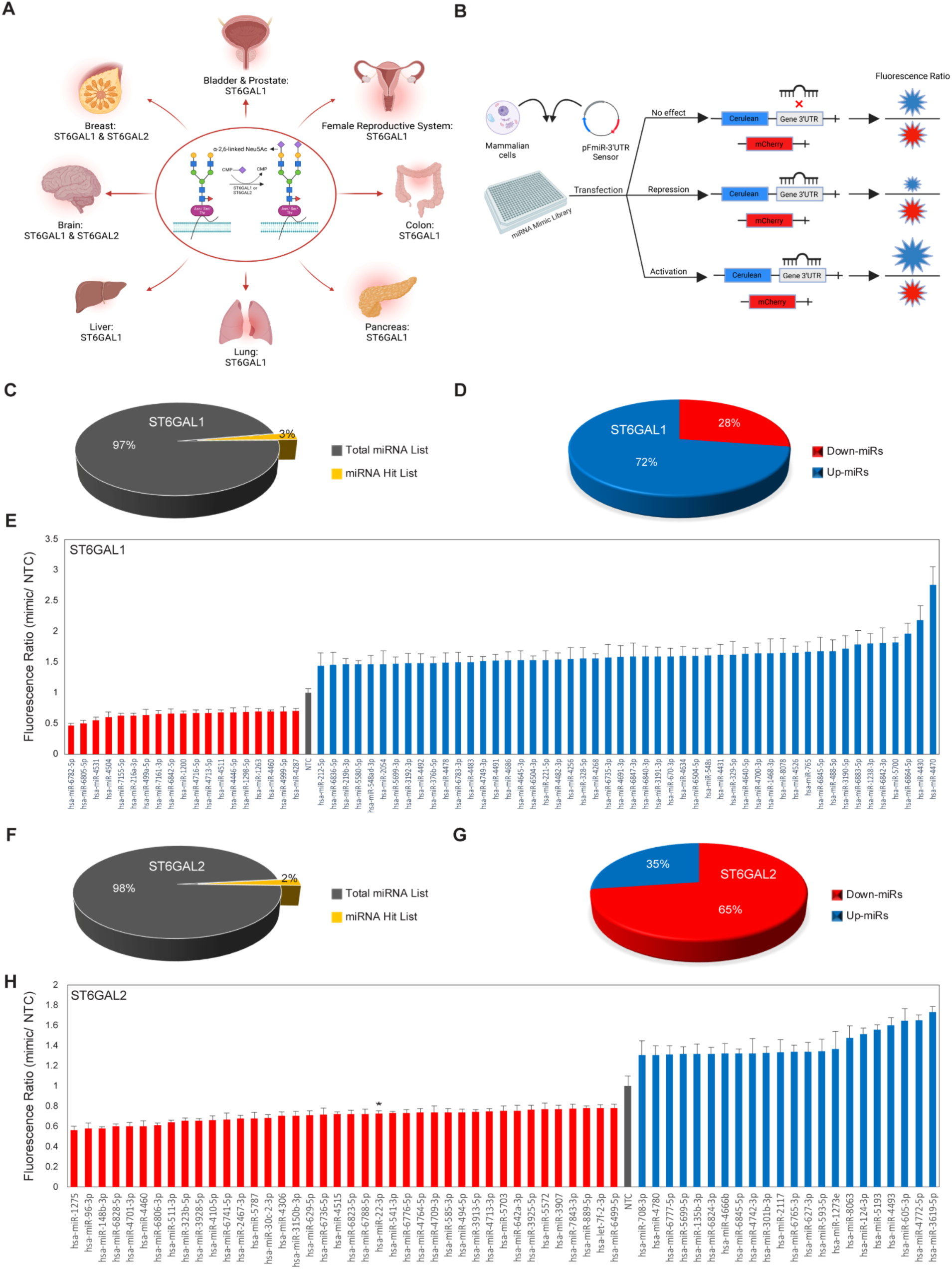
High-throughput analysis of miRNA regulation for ST6GAL1 and ST6GAL2 reveals different primary roles for miRNA. (**A**) Expression of ST6GAL1 and ST6GAL2. (**B**) miRFluR workflow. (**C** and **D**) Pie charts for ST6GAL1 representing percent (**C)** hits and (**D**) up- and down-miRs. (**E**) Bar graph of miRNA hits for ST6GAL1. Data is normalized to non-targeting control (NTC). Error bars represent propagated error. (**F** and **G**) Pie charts for ST6GAL2 as in C and D. (**H**) Bar graph for ST6GAL2 as in E. Star represents known hit (*16*).

Herein we use a high-throughput assay, miRFluR, to study miRNA regulation of α-2,6-sialylation (*9*). miRFluR uses a fluorescent sensor to map regulation of target proteins via miRNA: 3’-UTR interactions (**Fig. 1B**). We tested 2,601 miRNA mimics against ST6GAL1 or ST6GAL2. Our analysis identified 69 miRNA hits for ST6GAL1 and 62 for ST6GAL2, with no overlap observed (**Fig.1, C-G, Figs. S1-3**). Unexpectedly, the majority of miRNA regulating ST6GAL1 were found have upregulatory interactions (>1.4-fold, up-miRs), while those regulating ST6GAL2 were primarily downregulatory (down-miRs), indicating the predominant mode of regulation of miRNA may be target-dependent.

We validated our findings for a subset of hits from our miRFluR assays (ST6GAL1: 4 down-miRs, 8 up-miRs; ST6GAL2: 3 down-miRs, 3 up-miRs). For ST6GAL1, we tested four cancer cell lines: A549 (lung, **Fig 2** and **Figs S4**), PANC1 (pancreatic, **Figs. S5 and S6**), HT-29 (colon, **Fig S7**,**A-D**) and OVCAR3 (ovarian, **Fig. S7, E-H**). Hits for ST6GAL2 were studied in A549 (**Fig. 3 and Fig. S8**) and HT-29 **(Fig. S9**). Consistent with previous work, our assay accurately identified regulation of the endogenous proteins by miRNA (*9*). Overall, up- and down-miRs had the anticipated impact on ST6GAL1 and ST6GAL2 protein levels. In contrast, a more varied response was observed in mRNA, consistent with work showing discrepancies between mRNA and protein levels (*7, 9, 17*). We tested the impact of up- and down-miRs targeting ST6GAL1 on α-2,6-sialylation using *Sambucus nigra* lectin (SNA, **Fig. 2E, Fig. S4, G and Fig. S5, E**). Our results were in line with the effects of miRNA on protein expression, with up-miRs increasing and down-miRs decreasing α-2,6-sialic acid levels.

**Fig. 2.**
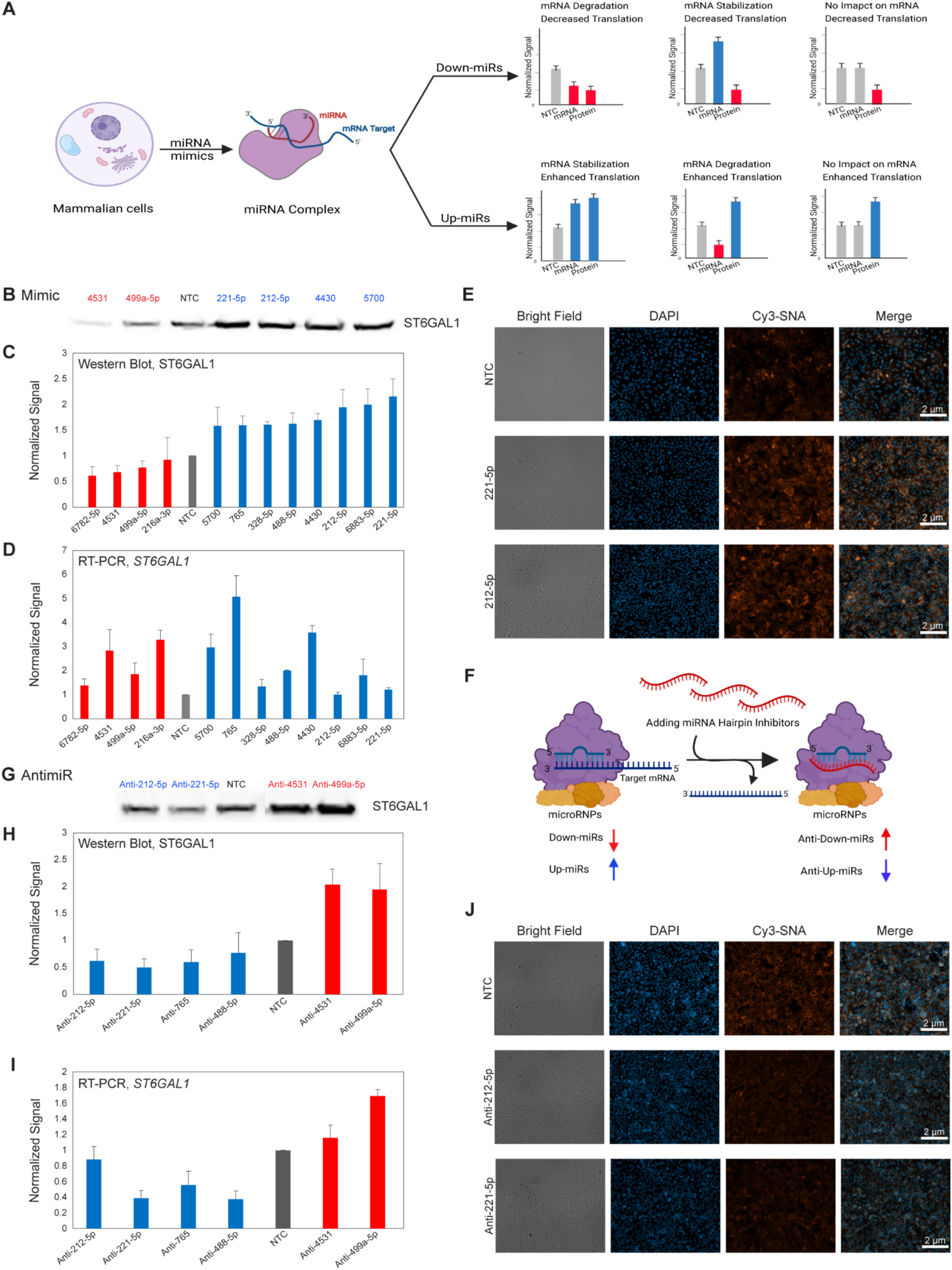
miRNAs regulate α-2,6-sialylation via controlling ST6GAL1 expression at both the mRNA and protein levels. (**A**) Scheme of possible outcomes. (**B**) Western blot of ST6GAL1. A549 cells were transfected with miRNA mimics or non-targeting control (NTC, 50 nM, 48 h). (**C**) Quantitation of experiments performed as in B. ST6GAL1 expression was normalized by Ponceau and divided by the normalized signal from NTC. miRNAs indicated in figure (blue: up-miR, red: down-miR). (**D**) RT-qPCR analysis for samples as in B. Data was normalized to GAPDH and to NTC. (**E**) SNA staining of up-miR treated cells as in B (NTC, miR-221-5p or miR-212-5p). (**F**) Schematic representation of antimiRs. (**G**) Western blot of ST6GAL1 for antimiRs. A549 cells were transfected with antimiRs or NTC (50 nM, 48 h). (**H**) Quantitation of experiments performed as in G and normalized as in C (n=3,). (**I**) RT-qPCR for experiments as in G. Samples were normalized to GAPDH and NTC. (**J**) SNA staining of cells treated as in G with NTC, anti-miR-221-5p and anti-miR-212-5p. Additional data including data in other cell lines are shown in Figs. S4-7. All experiments were performed in biological triplicate. Errors shown are standard deviations.

**Fig. 3.**
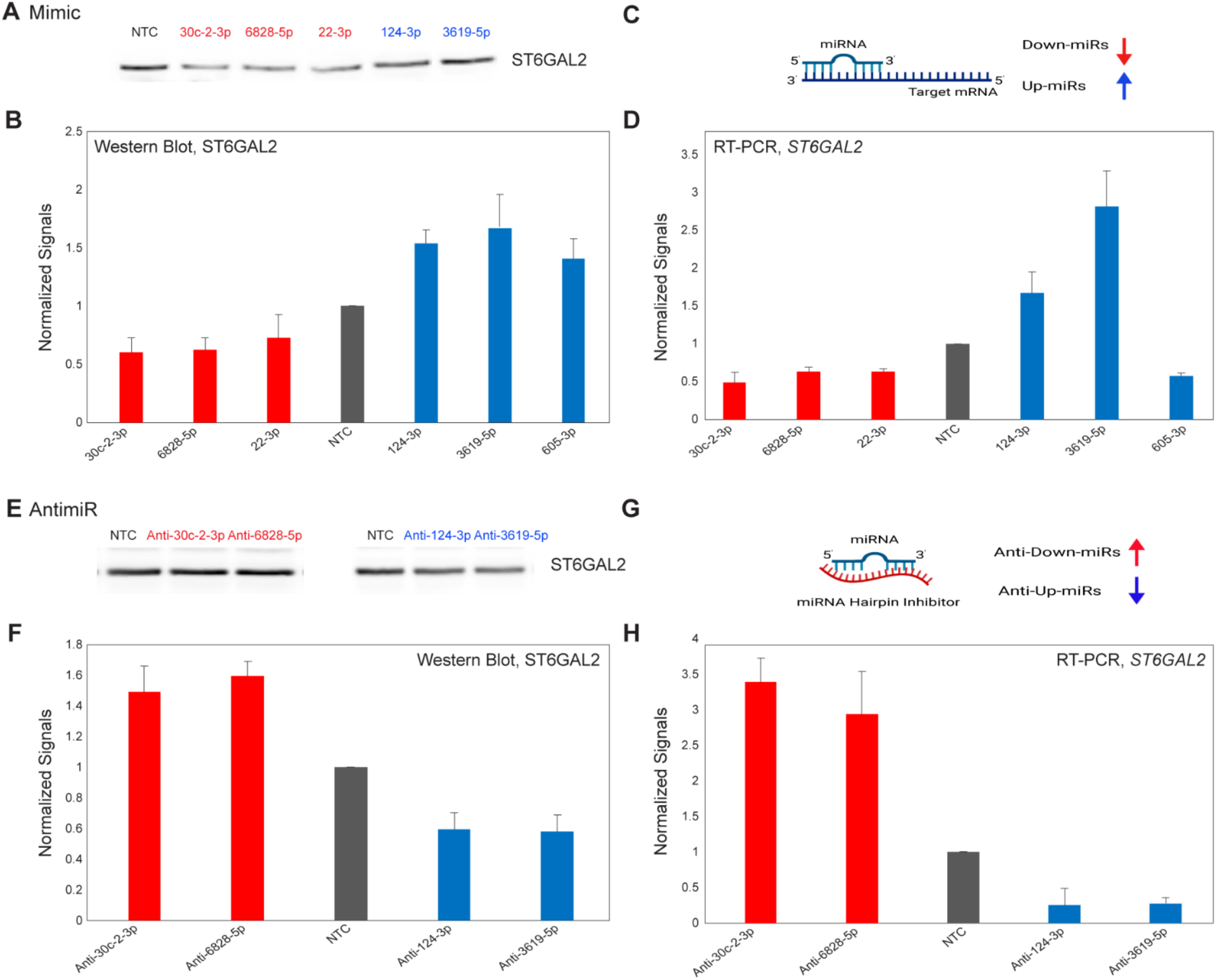
miRNAs regulate ST6GAL2 expression at mRNA and protein levels. (**A**) Western blot of ST6GAL2. A549 cells were treated with miRNAs as in Fig. 2, B. (**B**) Quantitation of Western blot analysis. (**C**) Scheme of miRNA regulation. (**D**) RT-qPCR analysis corresponding to B. (**E**) Western blot analysis of ST6GAL2. A549 cells were treated with antimiRs as in Fig.2, G. (**F**) Quantitative Western blot analysis. (**G**) Scheme of anti-miR regulation. (**H**) RT-qPCR corresponding to F. Additional data is shown in Figs. S8-9. All experiments were performed in biological triplicate. Errors shown are standard deviations.

We used antimiRs, which bind to endogenous miRNA and inhibit their actions, to further validate our findings (ST6GAL1: **Fig. 2F-J, Figs. S4, C-D and H, S6**. ST6GAL2: **Fig. 3, E-H, Fig. S8, C-D**). Our observations were in line with the expectation that anti-up-miRs downregulate protein expression and anti-down-miRs upregulate protein expression. For example, inhibition of miR-221-5p or −212-5p, up-miRs for ST6GAL1, caused downregulation of the endogenous protein in A549 and PANC1 (**Fig. 2, G-H, Fig. S6, A-C**). We observed concomitant loss of α-2,6-sialic acid (**Fig 2J**). This supports a function for these miRNAs in maintenance of sialyltransferase and sialylation levels through upregulation.

miRNA-mediated upregulation of protein expression in dividing cells has generally been attributed to either a direct impact of miRNA on promoters and enhancers, competition between miRNA, or indirect effects (*18, 19*). Our identification of up-miRs via miRFluR precludes that this regulation is through miRNA modulation of gene promoter or enhancer elements. Targetscan predicted sites for 47% of down-miRs and 38% of up-miRs regulating ST6GAL1, and RNAhybrid identified sites for all miRNA hits (**Fig. 4, A, Fig. S10**) (*20, 21*). Mapping of the predicted sites on ST6GAL1 showed that the majority of up-miR sites do not overlap with down-miRs (**Fig. S10**), arguing that the observed upregulation is not predominantly via miRNA competition. To test whether up-miRs act via direct base-pairing, we identified potential binding sites for miR-212-5p and miR-221-5p and mutated these sites in our sensors (**Fig. 4, B**). For both miRNA, mutation of one of the two potential sites caused a significant loss of upregulation, validating this as a direct effect (**Fig. 4, C**). The validated site for miR-221-5p was predicted by Targetscan and has a 6-mer seed region. The site for miR-212-5p was non-canonical with 10 of 12 base pairs matching at the 5’ end of the miRNA. In contrast to the work of Steitz and coworkers, neither of these validated sites were AU-rich (*3*), arguing that this motif is not necessary for activation of protein expression by miRNA. Sites for both up-miRs were in the same region of the 3’-UTR, although their “seed” regions did not overlap (**Fig. S10, C**). A significant number of the potential up-miR sites are in close proximity or overlap other up-miR sites, indicating a possible impact of local 3’-UTR structure on up-regulation. Mutation of a down-miR site (miR-4531) gave the expected result (**Fig. 4, B-C**). Our data strongly supports a mechanism for protein upregulation through direct interactions between miRNA and the 3’-UTR of mRNA in proliferating cells.

**Fig. 4.**
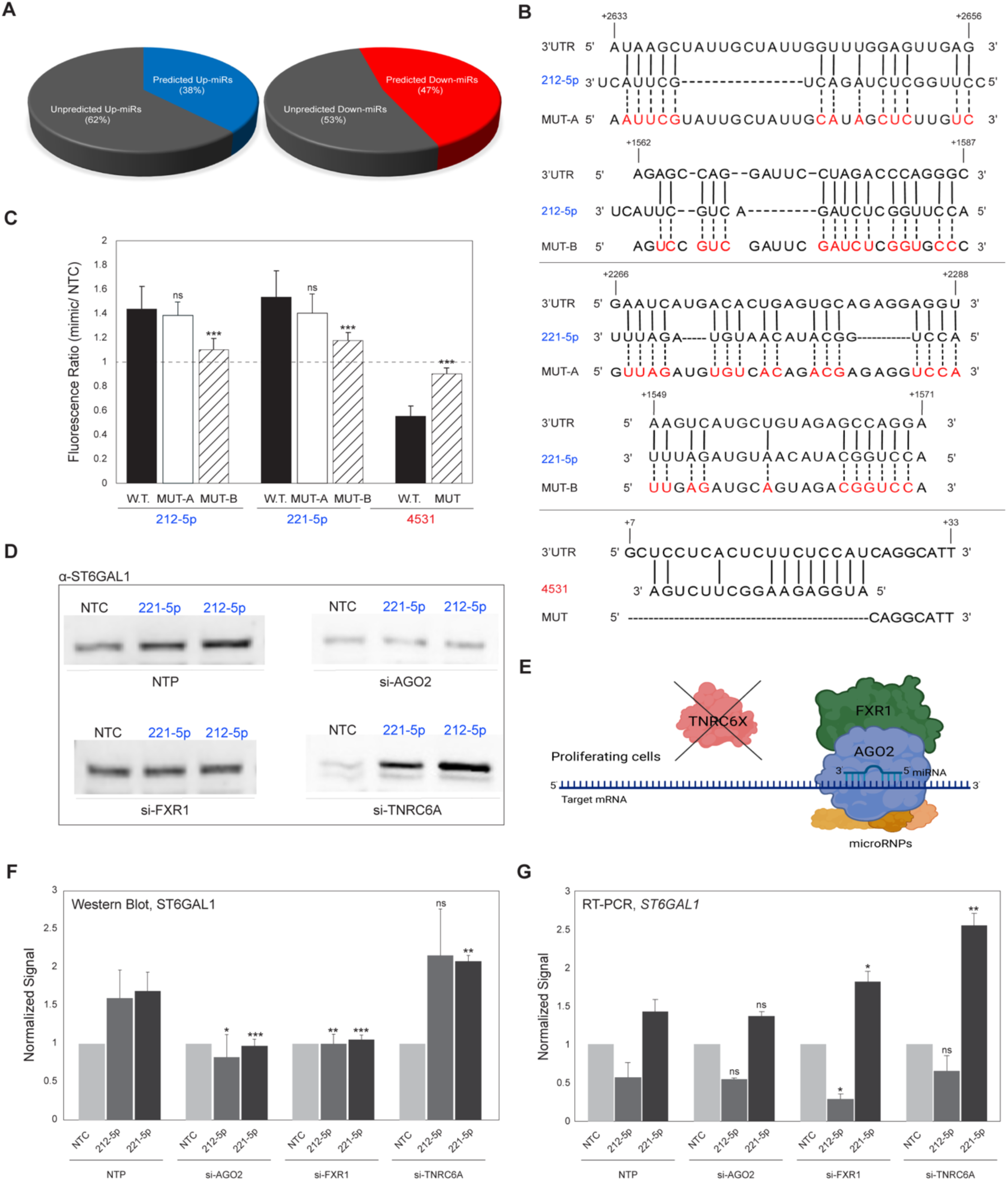
Upregulation of expression by miRNAs requires direct interaction with 3’UTR and utilizes FXR1 and AGO2. (**A**) Pie charts showing percentage of up-miRs (left, blue) and down-miRs (right, red) from Fig 1, E predicted by Targetscan for ST6GAL1. (**B**) Alignment of miRs (top: 212-5p, middle: 221-5p, bottom: 4531) with predicted 3’UTR sites and their corresponding mutants. (**C**) Bar graph of data from mutant miRFluR sensors as in B. Data was normalized to NTC in each sensor. For statistics data was compared to wildtype (W.T.) for each miRNA. (**D**) Representative Western blot of ST6GAL1. A549 cells were treated with pools of siRNA (non-targeting (NTP), si-AGO2, si-FXR1, si-TNRC6A, 48 h) prior to treatment with miRNA mimics (NTC, miR-212-5p, miR-221-5p, 48 h) and analysis. (**E**) Schematic representation of potential miRNA complex. (**F**) Quantitative Western blot analysis of experiment as in D. ST6GAL1 expression normalized as before. (**G**) RT-qPCR analysis for samples as in F. All samples are normalized to GAPDH and NTC. Additional data is shown in Figs. S8-9. All experiments were performed in biological triplicate. Errors are standard deviations. Standard t-test was used (*\* p < 0*.*05, ** < 0*.*01, *** <0*.*001*).

Steitz and coworkers were the first to propose a miRNA mediated upregulation pathway, in which miRNA that bound to AU-rich elements (AREs) could activate translation in senescent cells (*3*). This activation required the miRNA binding proteins Argonaute 2 (AGO-2) and Fragile-X-metal retardation related protein 1 (FXR1). In mitochondria, the absence of Trinucleotide repeat-containing gene 6A (TNRC6A, GW182) was noted as a critical requirement for miRNA-mediated activation (*5*). To test whether these proteins play a role in the observed upregulation of ST6GAL1 by up-miRs, we first used pooled siRNA to deplete the endogenous proteins in actively dividing A549 cells. Post-depletion, we transfected silenced cells (control (NTP), si-AGO2, si-FXR1, si-TNRC6A) with either non-targeting control (NTC) or individual up-miRs: miR-212-5p or miR-221-5p (**Fig. 4, D-G, Figs. S11-12**). Depletion of either AGO2 or FXR1 prevented protein upregulation by up-miRs, in line with Steitz’s work in senescent cells.

As previously observed, the changes in mRNA were not concordant with protein levels. In contrast, depletion of TNRC6A enhanced upregulation by miR-221-5p compared to the control (∼23% increase, *p* < 0.01). This is in line with an earlier proposal that TNRC6A inhibits upregulation (*5*). Similar results were seen with miR-212-5p, but they did not meet the statistical threshold. It should be noted that the pooled siRNA for TNRC6A did not strongly silence this protein (**Fig. S11, C**). Overall, our work supports a role for AGO2/FXR1 complexes in activation of translation in response to up-miRs in actively dividing cancer cells.

Currently, the common presumption is that in proliferating cells the direct impact of miRNA on protein expression is downregulatory. There have been a few cases, buried in the literature, contradicting this assumption (*9, 22, 23*), however this is not thought to be a major aspect of miRNA regulation. Our data for ST6GAL1, ST6GAL2 and previously B3GLCT (*9*) contradicts this, revealing that upregulatory interactions are more commonplace than previously thought. Consistent with this, high-throughput analysis of miRNA interactions for POT1, PTEN, MXI1 and other cancer-related genes also identified significant upregulatory interactions, but these were ignored as noise (*24*). In some cases (ST6GAL1), upregulation can be the major mode of regulation by miRNA, although these same miRNAs may have downregulatory activity for other genes. Like most glycogenes, both mRNA and protein expression of ST6GAL1 are on the lower end of expression levels in the cell. At this range of expression, noise becomes an increasing problem and miRNA are thought to be critical in correcting this (*25*). α-2,6-sialylation acts as an important regulator of a host of functions within mammals, including our adaptive immune system (*12*) and cell migration (*13*). Thus, it may make sense that regulation by miRNAs is in both directions to precisely control expression of critical low abundance proteins. Proteins in this class include glycosylation enzymes, GPCRs and most cell surface receptors. These proteins, which often act as initiators of amplified signals would be important to tightly regulate. The discovery of upregulation as a feature of miRNA control in proliferating cells expands our understanding of the miRNA regulatory landscape.

## Acknowledgments

The authors would like to acknowledge Dr. Eva Hernando (NYU Langone) for helpful insights and Dr. Matt Macauley (University of Alberta) for his generous gift of neuraminidase.

## Funding

This work was supported by the Canada Excellence Research Chair Program (CERC in Glycomics, L.K.M.).

## Author contributions

F.J.C. and L.K.M. conceived the project. F.J.C., D.M. and L.K.M. designed the experiments. F.J.C. and H.H.N. performed experiments. F.J.C. and L.K.M. did data analysis. F.J.C. and L.K.M. wrote the manuscript. All authors discussed results and edited the manuscript.

## Competing interests

Authors declare that they have no competing interests.

## Data and materials availability

All data are available in the main text or the supplementary materials.

## Supplementary Materials

Materials and Methods

Supplementary Text

Figs. S1 to S12

Tables S1

Data S1-Excel file

## Notes

### Competing Interest Statement

The authors have declared no competing interest.

