## Supplementary Information for "miRNA upregulate protein and glycan expression via direct activation in proliferating cells"

**This PDF file includes:**

Materials and Methods
Supplementary Text
Figs. S1 to S12
Table S1
Data S1

**Materials and Methods**

Cloning of pFmiR-ST6GAL1-3'-UTR and pFmiR-ST6GAL2-3'-UTR:

ST6GAL1 and ST6GAL2 3'-UTRs were cloned from genomic DNA (gDNA) extracted from HEK293T cell line using QIAquick gel extraction kit (catalog #: 28706) and primers shown in Table S1. The amplicons were cleaned up using the PCR clean-up kit (catalog #: KTS1115). The 3'-UTR fragments were cloned using the NheI and BamHI restriction sites downstream of Cerulean in the pFmiR-empty backbone (*9*) using standard ligation protocols (NEB) and verified by Sanger sequencing (Molecular Biology Services Unit, University of Alberta). Large-scale endotoxin free DNA preparations were made for sequence-verified constructs (pFmiR-ST6GAL1 and pFmiR-ST6GAL2) using QIAGEN maxi-prep (catalog #: 12362 and catalog #: 19048). Plasmid maps for pFmiR-ST6GAL1 and pFmiR-ST6GAL2 and the glycogenes' 3'-UTR sequences can be found in Fig. S1 and Fig. S2, respectively.

Cell Lines

All cells lines (HEK-293T, A549, PANC1, HT-29, OVCAR-3) were purchased directly from the American Type Culture Collection (ATCC) and cultured using suggested media (HT-29 & HEK-293T: Dulbecco's Modified Eagle Medium (DMEM), 10% FBS; A549: FK-12, 10%, FBS; PANC1: DMEM* (catalog # 30-2002), 10% FBS; OVCAR3: RPMI-1640, 20% FBS with 0.01 mg/mL bovine insulin) under standard conditions (5% $CO_2$, 37 °C). All cells used were below passage number 15.

miRFluR High-throughput Assay

The Human miRNA mimic library version 21 (MISSION, Sigma) was resuspended in ultrapure nuclease-free water (REF. #: 10977-015, Invitrogen) and aliquoted into black 384-well, clear optical bottom tissue-culture treated plates (Nunc). Each plate contained three replicate wells of each miRNA in that plate (1.8 pmol/well). In addition, each plate contained a minimum of 6 wells containing non-targeting control (NTC). To each well was added 20 ng of pFmiR-ST6GAL1 or pFmiR-ST6GAL2 plasmids in 5 µl Opti-MEM (Gibco) and 0.1 µl Lipofectamine™ 2000 (Life Technologies) in 5 µl Opti-MEM (Gibco). The solution was allowed to incubate at room temperature for 20 min. HEK293T cells (25 µl per well, 400 cells/µl in phenol red free DMEM with FBS 10%) were then added to the plate. Plates were incubated at 37°C, 5% $CO_2$. After 48 hours, the fluorescence signals of Cerulean (excitation: 433 nm; emission: 475 nm) and mCherry (excitation: 587 nm; emission: 610 nm) were measured using the clear bottom read option (SYNERGY H1, BioTek, Gen 5 software, version 3.08.01).

Data Analysis

We calculated the ratio of Cerulean fluorescence over mCherry fluorescence (Cer/mCh) for each well in each plate. For each miRNA, triplicate values of the ratios were averaged and the standard deviation (S.D.) obtained. We calculated % error of measurement for each miRNA (100 × S.D./mean). As a quality control measurement (QC), we removed any plates or miRNAs that had high errors in the measurement (median error ±2 S.D. across all plates) and/or a high median error of measurement for the plate (>15% for ST6GAL1 and >14% for ST6GAL2). After QC we obtained data for 2,161 miRNAs for ST6GAL1 and 2,166 miRNAs for ST6GAL2 out of 2601 total miRNAs screened. The Cer/mCh ratio for each miRNA was then normalized to the Cer/mCh ratio for the NTC *within that plate* and error was propagated. Data from all plates were then combined and z-scores calculated. A z-score of ±1.965, corresponding to a *two*-tailed *p*-value of 0.05, was used as a threshold for significance. Post-analysis we identified 69 miRNA hits for ST6GAL1 and 62 for ST6GAL2 (**see Figs. 1, S3 and Data S1**).

Western Blots for ST6GAL1 and ST6GAL2

Cells were seeded in six-well plates (80,000 cells/well) and cultured for 24 h in appropriate media. Cells were then washed with HBSS and transfected with miRNA mimics (50 nM mimic, Dharmacon, Horizon Discovery, 5 μL Lipofectamine 2000, Life Technologies in 250 μL OptiMEM ). The media was changed to standard media 12 hours post-transfection. Cells were then lysed at 48 h post-transfection in cold RIPA buffer supplemented with protease inhibitors. For Western blot analysis, 50 μg of protein was run on 10% gels (SDS-PAGE) and transferred to iBlot2 Transfer Stacks (nitrocellulose, Invitrogen, catalog number: IB23002) using the iBlot2 transfer device (Invitrogen). Blots were incubated with Ponceau S Solution (Boston BioProdcuts, catalog #ST-180) for 10 min and the total protein levels were imaged using protein gel mode (Azure 600, Azure Biosystems Inc.). Blots were then blocked with 5% (PANC1, HT-29. OVCAR3) or 10% (A549) non-fat dry milk in TBST buffer (TBS buffer plus 0.1% Tween 20) for 1.5 hours at 55 rpm on rocker (LSE platform rocker, Corning) at room temperature. For ST6GAL1 blots were incubated with rabbit α-human-ST6GAL1 1° antibody (1:900 in TBST with 10% non-fat dry milk, catalog #: 14355-1-AP, Proteintech). For ST6GAL2, rabbit α-human-ST6GAL2 1° antibody (1:900 in TBST with 10% non-fat dry milk, catalog #: 28367-1-AP, Proteintech) was used. After overnight incubation at 4 ℃, blots were washed 4× for 2 min each with 0.1% TBST buffer. After washing, a secondary antibody was added (α- rabbit IgG-HRP, 1: 10,000 in TBST with 10% non-fat dry milk, Abcam). After incubation for 1 h at room temperature with shaking (60 rpm), blots were washed 4× for 2 min each with 0.1% TBST buffer. The blots were then developed using Clarity and Clarity Max Western ECL substrate according to the manufacturer's instructions (Bio-Rad). Membranes were imaged chemiluminescent mode (Azure 600, Azure Biosystems Inc.). Western blot analysis was conducted for ST6GAL1 up-miRs in four cell lines (A549, PANC1, HT-29, OVCAR3; miRs: miR-328-5p, -488-5p, -221-5p, -6883-5p, -5700, -765, -212-5p, -4430) and for ST6GAL1 down-miRs in three cell lines (A549, PANC1, HT-29; miRs: miR-6782-5p, -499a-5p, -216a-3p, -4531). For ST6GAL2, both up- and down-miRs were tested in two cell lines (A549, HT-29; up-miRs: miR-3619-5p, -124-3p, -605-3p, down-miR: miR-30c-2-3p, -6828-5p, -22-3p). All analysis were done in biological triplicate.

The α-human-ST6GAL1 1° gave multiple bands in some cell lines. Therefore, we validated the antibody using the ON-TARGETplus siRNA reagent against ST6GAL1 in a smart pool format (Dharmacon, Horizon Discovery, CA) in PANC1 and A549 using the manufacturer's protocol (see Fig. S4).

RT-qPCR

Total RNA was isolated from cells treated as in Western blot experiments using TRIzol reagent (catalog #: 15596018, Invitrogen) according to the manufacturer's instructions. RNA concentrations were measured using NanoDrop, and high-quality isolated total RNA was reverse-transcribed to cDNA using Superscript III CellsDirect cDNA synthesis kit (catalog #: 18080300, Invitrogen). Reverse transcription quantitative PCR (RT-qPCR) was performed using the SYBR Green method and cycle threshold values (Ct) were obtained using an Applied Biosystem (ABI) 7500 Real-Time PCR machine and normalized to housekeeping gene GAPDH. The primer sequences used in RT-qPCR can be found in Table S1. All analysis was done in biological triplicate

SNA Staining Assay

Cells were seeded onto sterile 22 × 22 no. 1 coverslips placed into 35 mm dishes at a density of 5 × $10^4$ cells/ml in standard media. After 24h, cells were transfected with miRNA mimics as in the Western blot section. At 48 h post-transfection, cells were washed with PBS (3×, 2 mL) and fixed with 4 % paraformaldehyde for 15 min. Cells were again washed with PBS (3×, 2 mL). blocked using 10 % BSA in PBS for 1h in incubator (37 ℃, 5% $CO_2$) and Cy3-SNA was added (1:300 in 10 mM HEPES, 0.15 M NaCl, 0.1 mM $CaCl_2$, pH 7.5, Vector Laboratories, catalog # CL-1303). After 1 h in the incubator, coverslips were washed (PBS, 3×), and cells were counterstained with Hoechst 33342 (1 μg/mL in PBS, 15 min in incubator). The coverslips were then mounted onto slides with 60 µl of mounting media (90% glycerol in PBS) and imaged with a Zeiss fluorescent microscope (Camera: Axiocam 305 mono, software: ZEN 3.2 pro). Specificity of SNA staining was confirmed by using neuraminidase (gift of Dr. Matthew Macauley) prior to SNA staining. All analysis was done in biological triplicate

Endogenous miRNA Activity Validation Using miRNA Hairpin Inhibitor

miRIDIAN microRNA Hairpin Inhibitors (ST6GAL1: anti-miR-221-5p, anti-miR-212-5p, anti-miR-488-5p, anti-miR-765, anti-miR-499a-5p, anti-miR-4531; ST6GAL2: anti-miR-3619-5p, anti-miR-124-3p, anti-miR-6828-5p, anti-miR-30c-2-3p) and miRIDIAN microRNA Hairpin Inhibitor Negative Control (NTC) were purchased from Dharmacon (Horizon Discovery, Cambridge, UK). A549 cells were seeded and incubated as described for Western blot. A549 cells were transfected with anti-miRNAs, 50 nM using Lipofectamine™ 2000 transfection reagent in OptiMEM following the manufacturer's instructions (Life Technologies). After 12 h media was changed to standard culture media. 48 h post-transfection A549 cells were lysed and analyzed for ST6GAL1 and ST6GAL2 protein and mRNA levels as previously described. For ST6GAL1, anti-up-miRs (anti-miR-221-5p, anti-miR-212-5p, anti-miR-488-5p, anti-miR-765) were also tested in the PANC1 cell line. All analysis was done in biological triplicate.

Multi-Site Mutagenesis in 3'-UTR of ST6GAL1 pFmiR Sensor

The 3'-UTR sequence of ST6GAL1 and the three miRNA sequences (miR-221-5p, miR-212-5p, miR-4531) were analyzed with the RNAhybrid tool which calculates a minimal free energy hybridization of target RNA sequence and miRNA. The two stable predicted miRNA: mRNA interaction sites were selected for designing mutant pFmiR-sensors. Multiple mutation sites were designed and mutant sequences were ordered for synthesis from GenScript Biotech or Integrated DNA Technologies (IDT). Each synthesized mutant fragment (221-MUTA-gBlock, 221-MUTB-gBlock, 212-MUTA-gBlock, 212-MUTB-gBlock) was amplified by standard PCR machine (Bio-Rad), using the primer sequences found in Table S1. Amplicons were cleaned up using Monarch PCR & DNA cleanup kit (catalog #: T1030S, NEB). The NucleoSpin Gel and PCR Clean-up XS kit (REF. #: 740611.50) was used for DNA gel extraction when needed to exclude non-specific bands. The amplicons were ligated into the empty pFmiR plasmid after enzymatic digestion using a pair of restriction enzymes for each gBlock (221-MUTA: PasI, BamHI; 221-MUTB: NheI, PasI; 212-MUTA: SwaI, BamHI; 212-MUTB: NheI, PasI). Sequences for the mutant pFmiR-ST6GAL1 sensors were then verified by sequencing and used in the miRFluR assay as described previously. A minimum of 3-wells were transfected per sensor and the analysis was done in 2 independent experiments.

siRNA Knock Down of microRNPs: AGO2, FXR1, TNRC6A and Impact on Upregulation

ON-TARGETplus siRNA reagents against AGO2, FXR1, TNRC6A in a smart pool format and ON-TARGETplus Non-Targeting Control Pool (NTP) were purchased from Dharmacon (Horizon Discovery, CA). A549 cells were seeded in six-well plates (50,000 cells/well) and cultured for 24 h in appropriate media. Cells were then washed with HBSS and transfected with each of the siRNA pools (50 nM, NTP, AGO2, FXR1 or TNRC6A, Dharmacon, Horizon Discovery) with Lipofectamine™ RNAiMAX transfection reagent (catalog #: 13778150, Thermofisher) following the manufacturer's instructions. Media was changed 12 hours post-transfection. After 48 hours, cells were then transfected with miR-221-5p, miR-212-5p, or NTC as previously described. Cells were then harvested for Western blot and RT-qPCR analysis as previously described. The knockdown efficiency for the siRNA was tested by Western blot analysis using 1:1000 dilution of 1° antibodies targeting AGO2 (catalog #: 67934-1-Ig, Proteintech), FXR1 (catalog #: 12295S, Cell Signaling) and TNRC6A (GW182) (catalog #: ab114857, Abcam) in 10 % non-fat dry milk in TBST (Figs. S11-12). Blots were processed as for ST6GAL1/2.

**A**

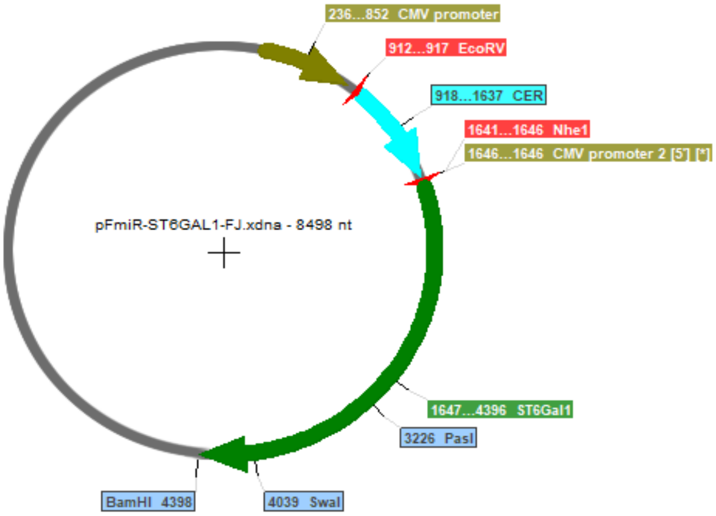


**B**

GCACAGGCTCCTCACTCTTCTCCATCAGGCATTAAATGAATGGTCTCTTGGCCACCCCAGCCTGGGAAGAACATTTTCCTGAACAATTCCAGCCT
GCTCCTTTTACTCTAGGGGCCTCTGTCAGCAAGACCATGGGGACTTCAAGAGCCTGTGGTCAGGAAATCAGGTCCAGCCTTCCCTGTAGCCAGAC
AGTTTATGAGCCCAGAGCCTCCTGCCACACACATGCACACATATCTAGCATTCTTTCCAGACAGCATCCTCCCCGCCTTCCACCTTGGTAGATGC
AAGGTCTATCTCTCCCATCAGGGCTGCCAAAGCTGGGCTTTGTTTTTCCCAGCAGAATGATGCCATTCTCACAAACCAATGCTCTATATTGCTTG
AAGTCTGCATCTAAATATTGATTTCACGTTTTAAAGAAATTCTCTTAAATTACAATTGTGCCCAATGCAGGGTGGCTCTGGGGGGCAAGTAGGTG
GTACAGGGGATTGGAAACATGCTCCGCGCCTCCAGAGAAAAGTTGCTCCCGAGGTCCATGCCCCTGGAACGTGTTCCTATCACTCTGGCTGGTTG
GGCTGGTCCTTAGACTGGGTGCTTATGATTAAAGGGTCTTGGTTAGCCCACTTTCCCTCTCCATGTGGAGATGGAAGGTAGAGAAGGATACAGTG
TCTATCCTCAAGTTGCTACGGTTCAGTGAGAGAGGCAGACATCTGAACAGGCAGGTAGGATTCAGTGTGCTCAGTGCACTGGGGATTTGGAGAGA
GATGGGCTTGCTCTCTCTGTGCACCCAGGAGGGCCACGCACTTAAAACTGTGTTTGTGGATCAGAGAAGGCTTTATAGCACAGGGGGCATTCAGA
TGAGTCTTAGAGGAAGAGAAGAAACATGGCAAGCAGATTACATCTGAGCCGTTTGAATTGTGTTTTTCTTTCTTCCCATGTTTATTTTCTAAGAT
CTACCTGAACTTAGAGACTCAAGATATTTTTTTAGGAAACCTCCTACCCATGTCTGAGGTAGCAAGTGCAGCCTCACGACAGATACCAGGCAATC
CAGAGCCACAAAACGTGATTCCTCCAGGCTCTGCCTGGCCTGACCCTGTCCTGTCAGCTGGGTTTACATACCAGTCCCATTCTTCCTTTTCAATA
CCTACCCCCAAATCTTCTCCTAACCACCATCTGTTTTTTTTTTTTTAAAGCATTTTTTGCTTTAAAAGCATCCTGACCCCAATTTCTTTGAGCTCA
CGGGCCTTTTGCTGAAGGTCTCTCAGGGTGTAGTGGTGTGGCTCTCTGGACTTAACGTCACTCTCAGAGGTCAGAACCTTGGAGATCAGAACTGA
TTCTCACCAGGTGTGAGAGGTGTGGTAGCAGATTGCAATGCTCTGCACCTCTTCCTTGCAAGTGAGCAACTTCAGGCTCTCTGGGCAGAGGCTGG
CCCACTGTAGTTTGCAGACATGCTCTCCAGATGGTTTTACTAAGTCCCCTCTCCCTGATAGGGAATCCTGCTGGACCAGCGCAGCCCTGGTGTGG
AGAGGTTAAAAGACTTGCACAGGATCACCAAGTCATGCTGTAGAGCCAGGATTCCTAGACCCAGGGCTCTGCACTCTCAAGGCTGGCCCCATGTG
CTCAAGGGGATCTAATGTTTGGGCTCCAAACTAACCATCTCGGAGCTGGGCTCCTCATTTACTGCCAAACCCTCAGCTTATGTAGCTAGAAAGGG
CCCTGGAGTGAGAAAGCCTGGATTTTCAAATTGATGCTCCCCTACTGACTAGCTGTGCCACTCTGGGCAAATGCTCTTCCTTGAGCCTGTTTCCA
CACCTGTAAAGTGGGGATGATGATCCTATCTCACTGCTTTTGTGAGGATTACAGGAAAGCACCTGTCCTGGCTCTGTACCTGGCACGTAGTAGGT
GCTCAGTTCATGCTGGTTTCCTTCCTGCCTTTAGTAGGGACCTGCTCTGTGCTCACACCTCGGCTGCATGCACCCTGCTGTGACGGAGGCTAGTG
TGGAAGAGGTCCTGTCCTCAGGGAATTAACTGTCTTATTGGGAGACAACAACTGTCCTCCTTGGAACACCCAAGAAACCATGCAAAGCAGTGGAC
AACACAGAACACGCCCTCCTCCTCGCTGCCTGCAGCTCCAATCTGATTCTGCTTGGGAATGGGCGGAGCACGTGGGCTGCTTAACTGCTGTATAG
GACAAGCCCCTTACCCCTCTCTGGGCCCATGAATTCCTGGCTTGGTTTATGTTCTGATTTGACACACTGATTTTAATCTTCGAATCATGACACTG
AGTGCAGAGGAGGTGGCATTCCGACAGCAGGACATACATGTTGGTGTGAAGACTGGGACGACACTGGGTAGAATCTAGTTTTTAATTATTATTAA
TATAAAGGATCAAATTAATTTAAATATGATTCTGAAGTCTACAGAACTTTTAGTTCTGTGCTGTCTATGTGGACACTTTGGTAAAATGCAAATTA
TGATATGGACGTTATCATTGGTCTGGTGAGATGTTTCATATTTGTGACAGTTAATTTAAAAATTATGAGTTAATGCTGCCTGTGTCTATGGGGTT
CTGTCTTCTTTGATAGCCATCTATTCATCTGGATCATGGGACCCTCTCTAATCCTTCCACCAATCAAATAAGCTATTGCTATTGGTTTGGAGTTG
AGATATCAGTCTCGGAAACTTCTGAAAAATGCTAATAATTACCCAAGGATTATGTCAAATTTTAAAATAAATGTGTGTGTGTTTCTTTAA

**Fig. S1.**
**pFmiR-ST6GAL1.** (A) ST6GAL1-pFmiR plasmid map. (B) ST6GAL1 3'-UTR sequence.

**A**

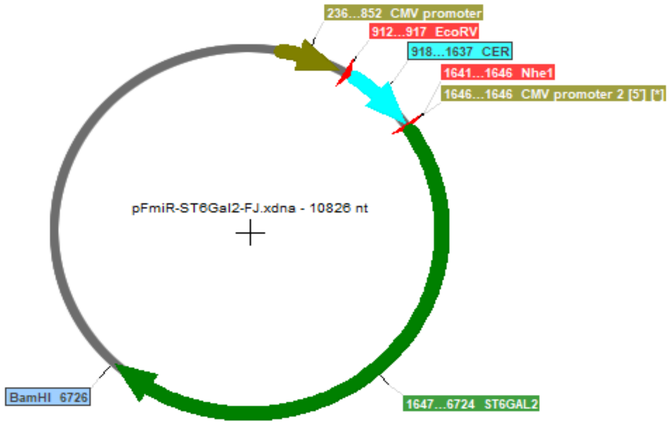


**B**

AAAGGGTTTCTTGGGAATCAATGTGCAATAAGGTACTACTGTTGTACTCAAAGTCACAAAAGAATACTTGAAGATAGATTTTAGACAGTGTAGTTTTG
AGTCTTTAAACAGTAATTTAGTGGTGGTCATGATAATCCTTCTCTTTCTAAGCCATTGAGTATGTATTACTGCTTTGAACATAGAAATGGTTTTTTTA
AAAAAATAAGGCTCAAGAAAGATGCAAAACCAAGGATAGCTAGAAAGAAAACACATGTATGACCATGGGTATGATACCTCCAGAAATGTTAACAACTT
ATTCTTCTGCCATGAGTACCCCTCATCAGGGTTGGTTTCAATGACAGGTTTTGGTGATGTTCCTGACCATATGAAGTGGTTTATGTTTAGAAACATTC
AAATTGAGGGACATCATTTACAGCATCGAGTGTGTCAGTTATACATGCATTCATCATTAAATATCCTAGGTTTTATGACAGACATTGAAATTATCACT
CAGATGTGTTTAACAGCAAAAAATTCCCAGCACATCCTGGCAAAGGCTTTTATTTCCAACCGTTGCATTCTTCATCTCTGCCTCCCATTGCCCACTGA
ATGCTTTGCTTTCTGTGCATCAAGACAGAGTTCTAAAACCAGAAAACATCCATCTTGAAAGGTCCAAGAAAAAGCCAAATTCCATATAGCACCACAGA
GGAGGCCCTTTGTTTCCCTGCAGTATTTCAACCAGGAACATTAGTATCACATCCACCAGGAGGCACATCACTGGCAAGTAACATAGAATCACAGGGCT
GCAGGTGCCCCTGGACACTCCCCGGTTCTGGCTTTTCTGCCTTGTACCTCCTCACCTGCAGATCTACCCTGCAGCTTTTGCATAGTCAGTGACACACT
TGACAATTGGCACAGCTGTGCTGGGGAAGACGGGAGACACCTTGCTCCTTCCAAACTAGAAGGAAAGATTTTGGAGCCATCACCCTGTGCTTCTAGGG
ATGCACGCCTGTCTTTATTATACAGGTTTCTTGCAGGTTTTTATCAGCCAGTGGGAACTCCACAGTGATGAGCACTGAATCTTCACTAAATCTTCCTA
CAGACATTTAGGAATCCAGTCCTTGCTTTTGCAGATATGTTTTATGTCAGCAGCATGACAAAGCCATATAATTAAATAATTTATGTGAAAGCATTGCT
TATTAACAAGGAAAATATGTAGATGCTTCAGAGGGAAAGGCAGCTAGAGGGAAATTTTCATCCCAATATGTTAGCTATATTCAGGGTTGATTTTTTCT
TTAATCTCAACTGTCACTGATACATTGAAAAATTTTTGCTATGAATATAGTGAAACAGAAACATTCTGTAATACCTTAATTAAACATGCTATTATTTC
AATGTATAACCAAAACATATCTCCATGGTACAGTTAAGACTTGTTGCAATGACAATAGCAGCTTAATTAAATGCGTTCTCCCTTGCAAAATGTATGAA
TGTCAGGAGAGTTTAGTGGAAAAGTTTTCTTTGTTTCAGGATTGGTTTCTTCTAGACTGTTCTCAGGACTACTTTCTTGAGCAAGAATGGCTTACAGA
GACTGAAGGCTTCTGTTTGGTTAGTCTATTCAGCAGATATGGAGCTGGAAAACAAGACATCTCCTATAGCTTGCTATAATAGTGAAGGAGAAATTGGA
AAGTTTTAATCACAATAATATCCAATAAAAGCCCATTTTGAGAGTTTTGTTATTAAGTCAATGAGAAGGGATAGGTAACTTTGTCCTATATATTAACT
TATGTCTAAATAGACATGGTTAAGCCATGTTCCTTTCTAATTTCTTTTCTTTTCTTTTTTTTTTTTTTTTTGAGACAGTCTCTCGCTCTGTCGCCCA
GGCTAGAGTGCAGTGGTGCGATCTTGGTTCACTGCCAGCTCTGCCTCCCACGTTCACACCATTCTCCTGCCTCAGCCTCCTGAGTAGCTGGGGACTAC
AGGCACCCGCCACCACGCCCAGCTAATTTTTTGTATTTTTAGTAGAGACAGGGTTTCACCGTGTTAGCCAGGGTGGTCTCGATCTCCTGACCTTGTGA
TCTGCCTGCCTCGGCCTCCCAAAGTGATGGGATTATAGGCGTGAGCCACCACCGCGCCCGGTCATTCCTTTCTAATTTCTTGAAAAAACTTTGTCTTG
TCCTTATGTTTACATGTGTGTAATAATTTCAGCAACTCTGGAATGTGGGTGAGGTGGCTGTGGTGGATGAATATTACTCCCTACCCTCCTTGGATCTT
TCTAGATCTTCCAGACCATTAAATGCCCCTGAAGCTCCTTTTATAGTTTTGCAAGTGGATTTACTTGTAGACATCCTTACTATCCAGCATAATCCCCC
TTGGCTGACATGAGAACTTGGTTAGAGGAGAATTTATTGGGGTCAGTGTAAATATTCAAGTTCAAAAGCCTCAGAACCCAACACTTTAAAGATTTTTA
AATGGAATTTTGTCGTAGTCATTGTACTTTTAAAATAGTACTTTCAGTTTCATTGAGGTTACAAGAAAGACTGAAGAGTACAAAGGAAATTTAGTTAC
AATTTTTGCATTTATTATGGCCTTAAAGACTCTTGGGAATCAGGCTCTGTTCAACAGTCAATAAATGTAATACACACATTAATATAGTACTGACTGGG
CGTTCTTCTATAAAAGTTTAAGTTCTTTGGCTTCTAAGAAAATGTTGGGCAGTTACAGTTAACTTGATGACTACTTCTAGTATTAGCCAAATGGGTCC
TAAGTGGCAGCATTTTGCCAGACATTGACTAAATCCTTACATGCCACATTTACTGAATTCCAATAAACATATAGTTCTTCTGTGGAGTAGTTAACAGT
AAGTGGCAGCTCACTGAATTTCTCCCAGGTTCTGTCTCTTCCAAAATCCTCCATCCTCTCTTATGAAGGGAAGCTGATAAAAAAAAATCCTCTTGGTG
CAAGCTGAAGCTGTTAAGGATTCCTTTTTTCACTCTCAGGACCAACAGACGTTCTTTTGTATTTCAAGATAATTTCACCTGTTGTTGTTGTTGTTGTT
GTTGTTGTTGTTGTTGTTGTTGTTGTTACAAAGAAACAAATTTTGAGCAGCAATGACAATTCAACCATATCTAGATTCTCCCTACCTTATTTCTCCAG
CCAGTCACACCAGGCAGACGGGTAACCTTTAACAGACCTGGTGTTCTCATGGAATACCGATGCTCGTCCCAGGGAGCTGACCTTTCTCACTATGTCCT
TATAGAGCTGCTCTTACAGAGAACTTTGTGCTCAGGACCAAGGGATGGTCAATTAAAAGTTGCATGTAAGCATCAGGCTTGGATGGGCAAGGGAGGGC
TTTTATACATCTTTGAACGACCAGGTCTTTCCTTGTCTGCCTGAAGGCAGAGCTTCGAGAAACAGCACTTCCCATGTGTCTGGTCCTCTTGGTCTCTC
AGCCAGACACTTAATCAATCCTGGACTCTCCAGTAGTCCTCAGTCATTGTAGCCATAGCATCTGCAAAAAAAGAAATGCTAAAATAATCATTAGCGTT
TTCAAAATTAGGAAATCAAAGTAGCTGGGTCACTCTCTTACATGTGTATGAGCCAGTTAATTCTGCTGGACAGATCATACTTACGCTGCTGATTGACT
CTTCTGAGCAGTAACACATCTCCTGAATTAATTAGGTAGGTCCCGTTTGGGTGCTTTAAGTGGGATCAGTCTTAGATGAGTGCCAGTATGTGATTGAA
TTTCATACGCACAAATGCCATGATTCAAGTTCTCTCTCCAGTTTTTCTTATAGATCAACACACAGTACATGTGGTCAGTTACTTGGTTGAATTTCTGC
TTAGCTGTGGTAAGAGACAGAGAAAGGGAGTGAGGTGGGACCAGGGGAATTATCAGAAAGACATGAGTGAACCTATTGGCCAGGTAGCTTCCAAGTAT
GTTCTATTGTATTGTCTGTTTTGATTCTGATAGTTTAGAACTCTTTCTGATAGTTTAGAACTCTTACCCAGTGGTTACATGACTGATCAAGGCAACTC
CATCTGGAAATTAATTGCTTGGGTAAAATGTCCCTGAGTTTCAACTTGTCTTTTATCTCTGGATATTCTCTTATGAGATTATTATGCATGTTTCTGTC
ACATGGGCATTATTTACACACTTGATAAATGAAAGGTATTTCTCTGTCCGTTCAAGGAAATAAAATGTAAGTTCTTGTTTTATTAAATGAATGAAGCC
TTTTGGTTAAAAAAAAAGTAATGTAATATGTTTCTGTTCAAACTTGAGCTAATGACATTTAAGGAATTAAAGACATTGATCTAGTTCTTAGTATGAAA
AGTGTACATGTAGATATAGTTATATAATTGTGTATATACATATCTACACACATGGTTGGTTTTATATTGCTTTTTCATCAATATTTTTGACTGAAACC
TTTACAATAAATTTTATTTTAATTATGTAACCACTAATATTTTGAGAGAAGGAAGCAAATGTTATATGTCCTTCTGTTAAAAGAAAAAAAATTCAGTC
CTCCGCCAACCGCTTCCCACACACTCATACTCTGCACATCTTGGCTACATTAAAATGGACTGACAGAATTATTTAAAAGGCACTATGAAGGTAGAACA
CTACCTTAATATTTTATCCTAGTCAGAAGCCTGATTTAACACTGAACTTTGAAGGGAAGCTTAGCTGTAGCTCATGGCTTTCAAAGATTAGAAGAAAC
ATTTACTGGTTCTATGGAAAATAGTCTTCTTGCTAGGTAGATTGGAAACAAAGAAATTGCATTGGTGGTGTTAAAATCTAACATTTATAACAAATCCT
GTTTTATTCTAGATGAAATACTTAGACAATTCTTCCAGTGGAAACCAGTGAACTTTTTTAACAGCTTTGGGAATTCTTTTGTATGATAATTTTTAAAA
TGAAATGTGTACTCTTCAAACTAATTAAAGCTTGAAGTTTTCCAGGAGTCGCATTTTAAAAGTGTTTGTACACATTTTAACACTTGGATATTTTCTAT
TTTGGTTTTGTATATTTTTATGTATACACAATGTACAGACTTTTTTCTTGTAAATAAAACATGTTTTCATTTGTCTAGAA

**Fig. S2.**
**pFmiR-ST6GAL2.** (**A**) ST6GAL2-pFmiR plasmid map. (**B**) ST6GAL2 3'-UTR sequence.

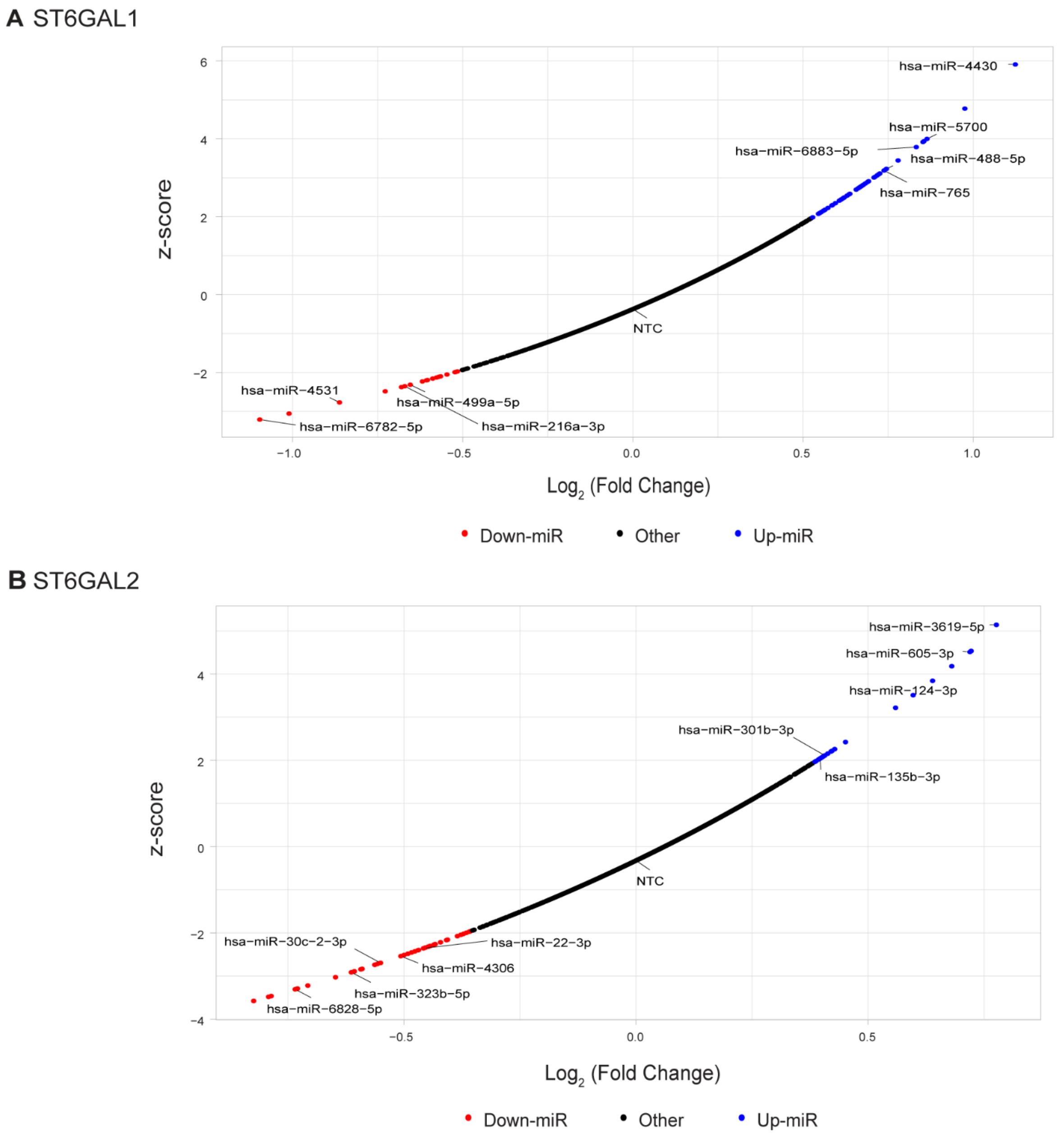


**Fig. S3.**

**Scatter plots of miRFluR assay showing up- and down-miRs.** (**A**) Data for ST6GAL1. (**B**) Data for ST6GAL2. miRNA in the 95% confidence interval are colored (down-miRs: red, up-miRs: blue).

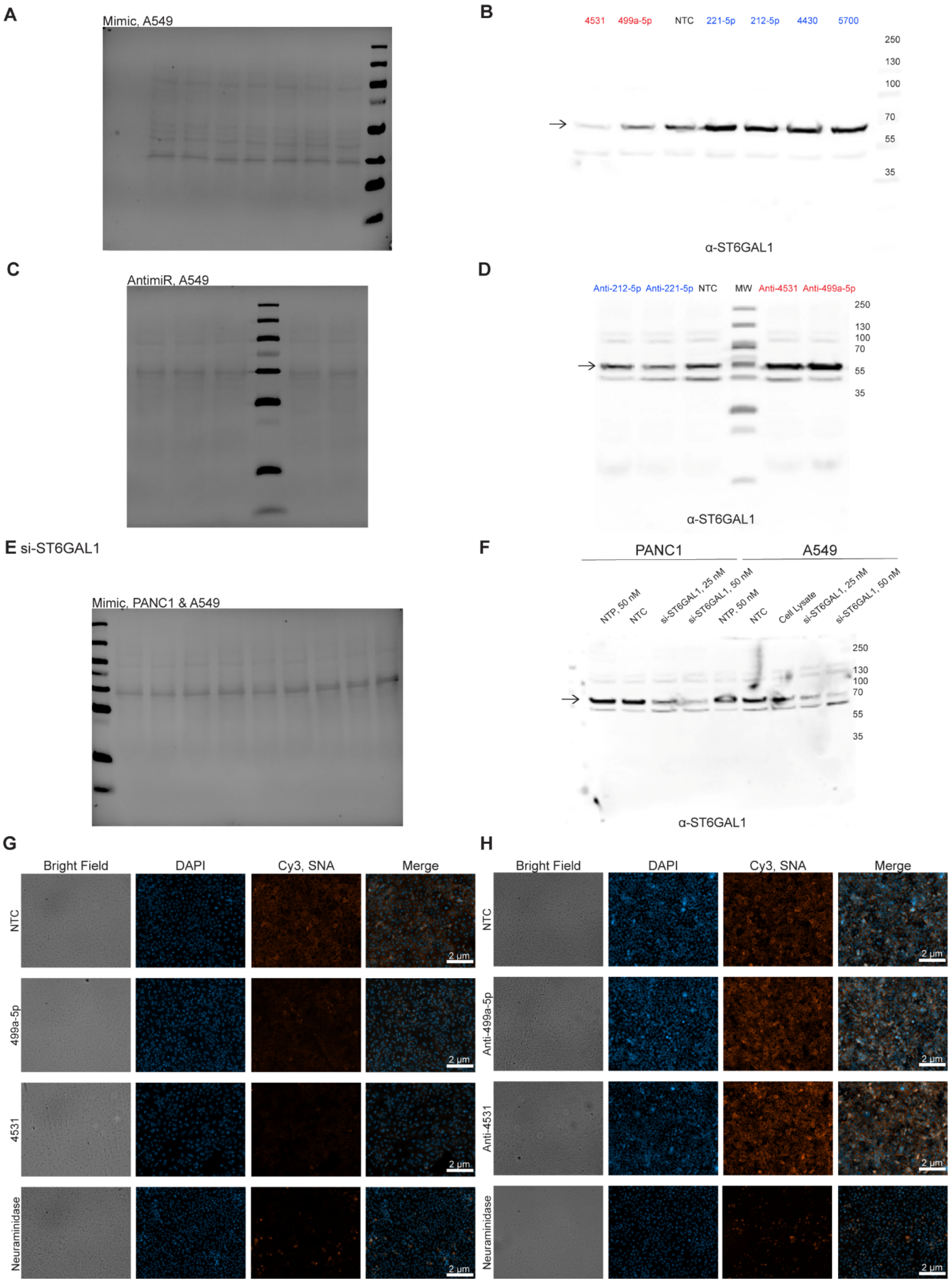
A
Mimic, A549
B
4531 499a-5p NTC 221-5p 212-5p 4430 5700
250
130
100
70
55
35
α-ST6GAL1
C
AntimiR, A549
D
Anti-212-5p Anti-221-5p NTC MW Anti-4531 Anti-499a-5p
250
130
100
70
55
35
α-ST6GAL1
E si-ST6GAL1
Mimic, PANC1 & A549
F
PANC1
A549
NTP, 50 nM
NTC
si-ST6GAL1, 25 nM
si-ST6GAL1, 50 nM
NTP, 50 nM
NTC
Cell Lysate
si-ST6GAL1, 25 nM
si-ST6GAL1, 50 nM
250
130
100
70
55
35
α-ST6GAL1
G
Bright Field
DAPI
Cy3, SNA
Merge
NTC
499a-5p
4531
Neuraminidase
2 µm
H
Bright Field
DAPI
Cy3, SNA
Merge
NTC
Anti-499a-5p
Anti-4531
Neuraminidase
2 µm

**Fig. S4.**

**miRNA up- and downregulate ST6GAL1 protein expression and α-2,6-sialylation activity in A549 human lung carcinoma cell line.** (**A**) Ponceau of Western blot shown in Fig. 2B. (**B**) Western blot shown in Fig. 2B. (**C**) Ponceau of Western blot shown in Fig. 2G (**D**) Western blot shown in Fig. 2G. (**E**) Ponceau of Western blot shown in F. (**F**) Validation of ST6GAL1 antibody. Pooled siRNA against ST6GAL1 were transfected into A549 and PANC-1 using standard protocols. In all ST6GAL1 Western blots arrow indicates the ~ 57 kDa ST6GAL1 protein validated by this knockdown. (**G**) SNA staining of A549 cells 48 h post-transfection with NTC or down miRs (miR-499a-5p, miR-4531). Neuraminidase treated cells are shown as a negative control. Our data shows ST6GAL1 protein downregulation by miRNAs correlates with decreased α-2,6-sialylation. (**H**) SNA staining of A549 cells 48 h post- transfection with NTC or anti-down-miRs (anti-miR-499a-5p, anti-miR-4531). Images are representative of n=3 experiments. Our data shows downregulation of ST6GAL1 by anti-miRs correlates with increased α-2,6-sialylation activity.

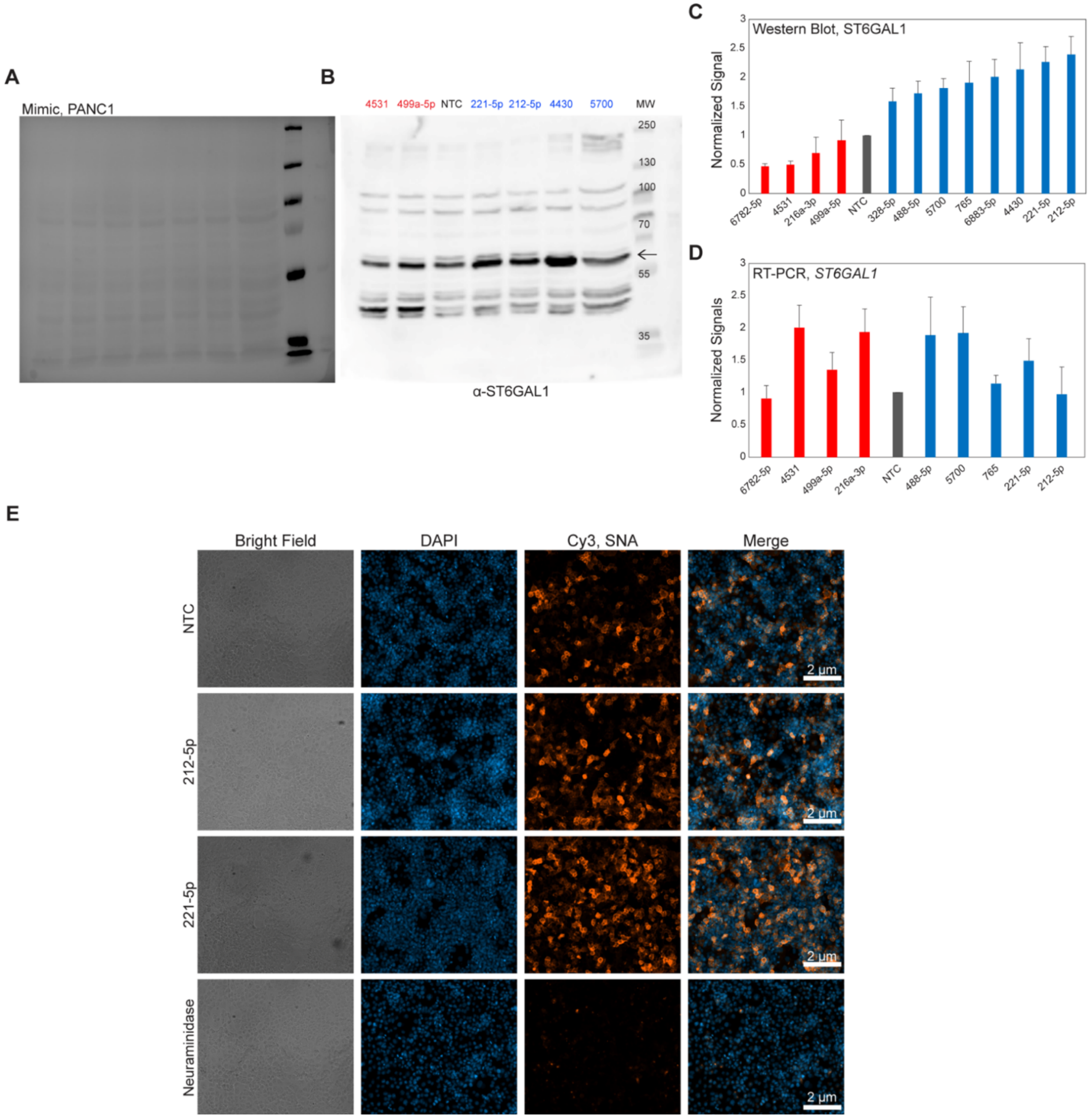


**Fig. S5.**

**miRNA up- and downregulate ST6GAL1 protein expression and α-2,6-sialylation activity in PANC1 pancreatic ductal carcinoma cell line.** (**A**) Ponceau of Western blot shown in B. (**B**) Western blot analysis of ST6GAL1 in PANC1 transfected with representative 50 nM miR mimics or NTC, 48 h post-transfection (arrow indicates validated ST6GAL1 band). (**C**) Quantitation of Western blot analysis illustrating up and downregulation of ST6GAL1 by individual miR mimics (up-miRs: miR-221-5p, 212-5p, -4430, -5700, -765, -328-5p, -6883-5p, -488-5p; down-miRs: miR-6782-5p, -499a-5p, -216a-3p, -4531; n=3). ST6GAL1 expression was normalized to total protein levels from Ponceau staining and set over normalized NTC for each blot. (**D**) RT-qPCR analysis for increased and decreased expression of ST6GAL1 mRNA by individual miR mimics. All samples are normalized to GAPDH as an endogenous housekeeping gene and then to NTC (n=3). (**E**) SNA staining of PANC1 cells 48 h post-transfection with NTC, up-miRs (miR-212-5p, miR-221-5p). Neuraminidase treatment are shown as a negative control. Data shown is representative of n=3 experiments.

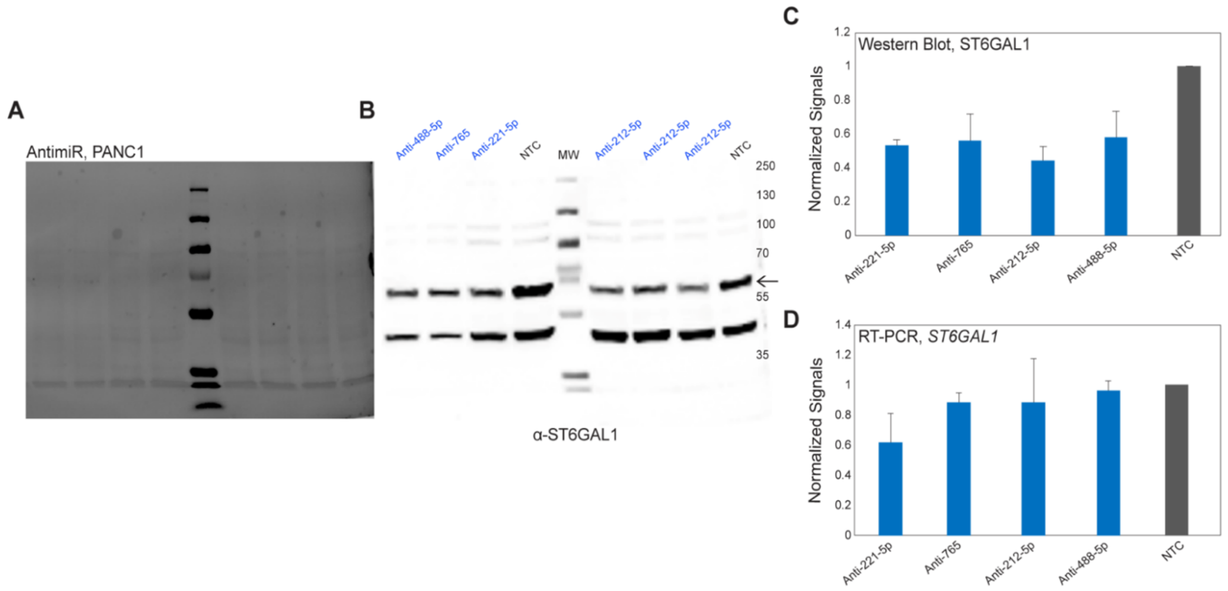


**Fig. S6.**

**Inhibiting endogenous up-miRs lowers ST6GAL1 protein expression levels in PANC1.** (**A**) Ponceau of Western blot shown in B. (**B**) Western blot analysis of ST6GAL1 in PANC1 transfected with 50 nM anti-up-miRs (anti-221-5p, -212-5p, -488-5p, -765) or NTC, 48 h post-transfection (arrow indicates validated ST6GAL1 band). (**C**) Quantitative Western blot analysis illustrating downregulation of ST6GAL1 protein by individual anti-up-miR (n=3). ST6GAL1 expression was normalized to total protein (Ponceau staining) and divided by normalized NTC for each blot. (**D**) RT-qPCR analysis for decreased expression of ST6GAL1 mRNA by individual anti-up-miR. All samples are normalized to GAPDH as an endogenous housekeeping gene and then to NTC (n=3).

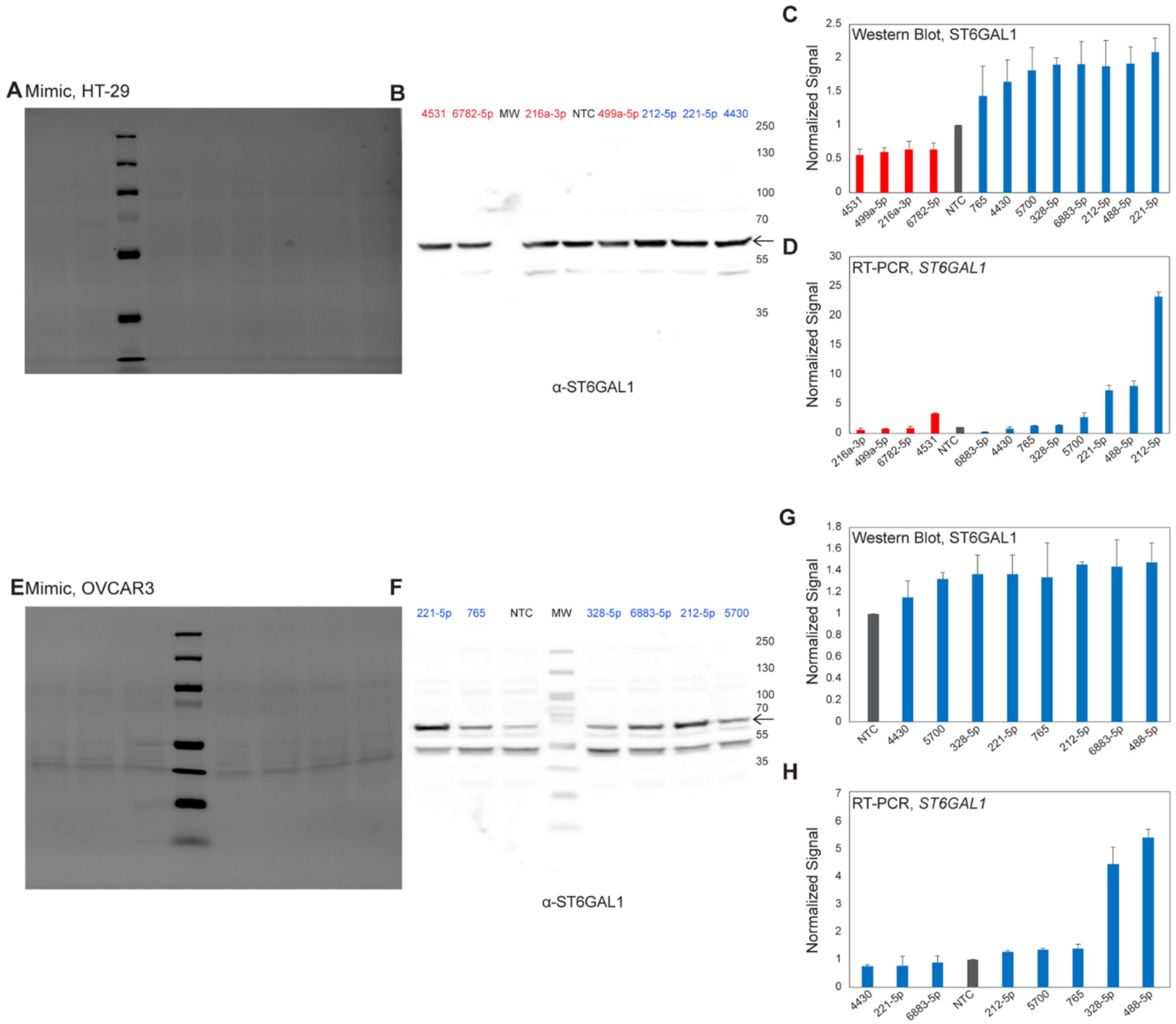


**Fig. S7.**

**miRNA up- and downregulate ST6GAL1 protein expression and α-2,6-sialylation activity in HT-29 human colorectal adenocarcinoma cell line and OVCAR3 high-grade serous ovarian adenocarcinoma cell line.** (**A**) Ponceau of Western blot shown in B. (**B**) Western blot analysis of ST6GAL1 in HT-29 transfected with 50 nM miR mimics or NTC, 48 h post-transfection. A representative set is shown. (**C**) Quantitative Western blot analysis illustrating up and downregulation of ST6GAL1 by individual miR mimics (n=3). ST6GAL1 expression was normalized to total protein (Ponceau staining) and divided by normalized NTC for each blot. (**D**) RT-qPCR analysis of ST6GAL1 mRNA from cells treated as in C. All samples are normalized to GAPDH as an endogenous housekeeping gene and then to NTC (n=3). (**E**) Ponceau of Western blot shown in F. (**F**) Western blot analysis of ST6GAL1 in OVCAR3 transfected with 50 nM up-miR mimics or NTC, 48 h post-transfection. Representative miRNA are shown. (**G**) Quantitative Western blot analysis illustrating upregulation of ST6GAL1 by individual up-miR mimics (n=3). ST6GAL1 expression was normalized as in C. (**H**) RT-qPCR analysis of ST6GAL1 mRNA from cells treated as in G. All samples are normalized to GAPDH as an endogenous housekeeping gene and then to NTC (n=3). Arrows indicates validated ST6GAL1 band.

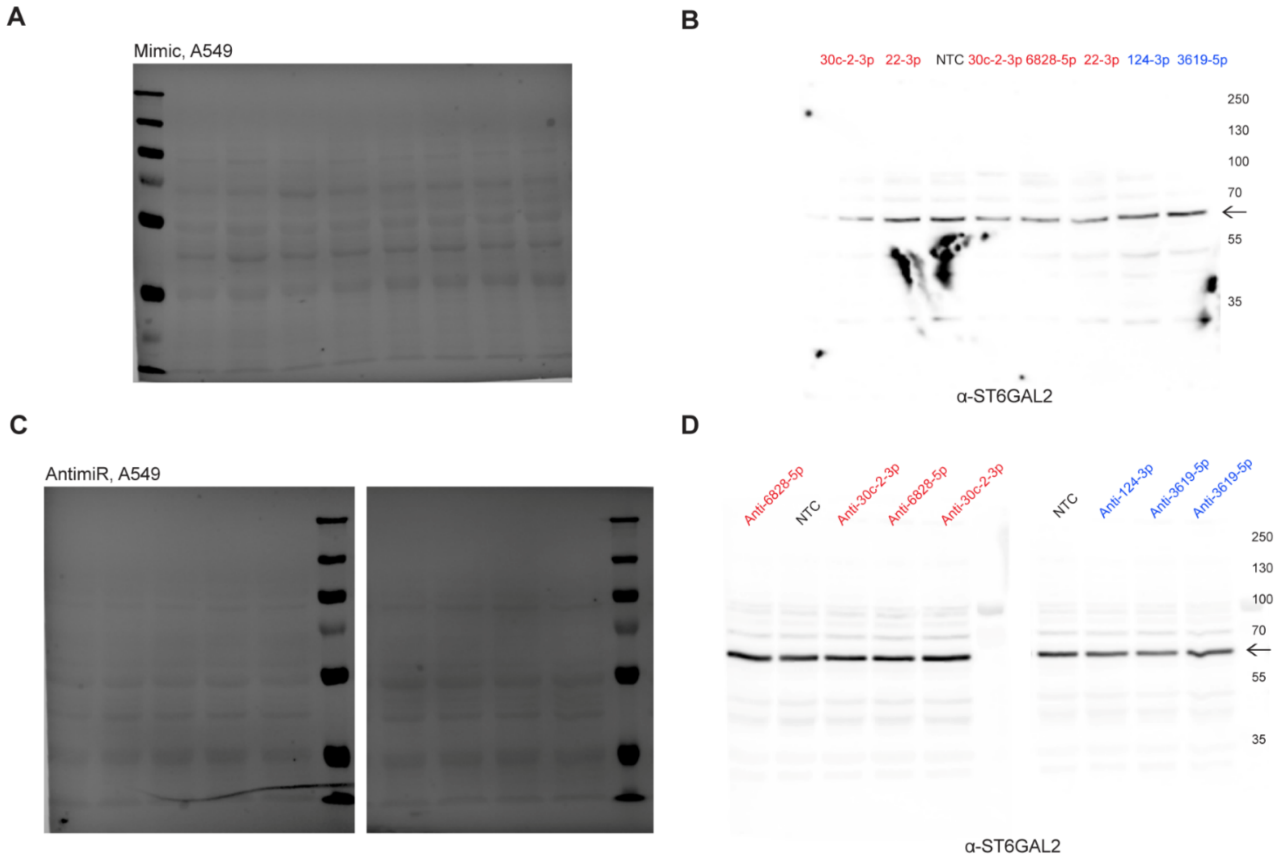


**Fig. S8.**
**miRNA up- and downregulate ST6GAL2 protein expression in A549 human lung carcinoma cell line.** (**A**) Ponceau of Western blot shown in Fig. 3A. (**B**) Western blot shown in Fig. 3A. (**C**) Ponceau of Western blot shown in Fig. 3E. (**D**) Western blot shown in Fig. 3E. Arrows indicates ST6GAL2 band (~ 60 kDa).

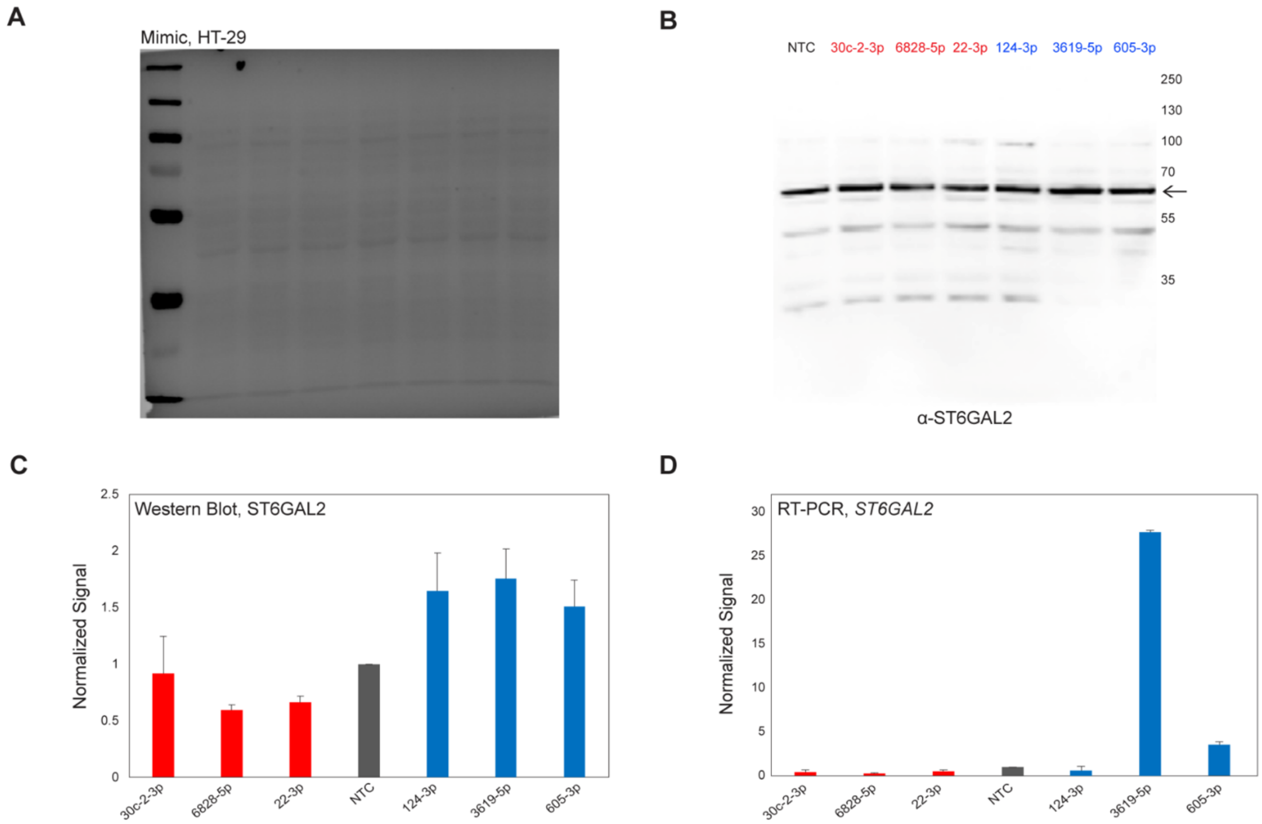


**Fig. S9.**
**miRNA up- and downregulate ST6GAL2 protein expression in HT-29 human colorectal adenocarcinoma cell line.** (**A**) Ponceau of Western blot shown in B. (**B**) Western blot analysis of ST6GAL2 in HT-29 transfected with 50 nM miR mimics or NTC, 48 h post-transfection. Select subset of miRNAs is shown. Arrow indicates ST6GAL2. (**C**) Quantitation of Western blot analysis illustrating up and downregulation of ST6GAL2 by individual miR mimics (up-miRs: miR-3619-5p, -124-3p, -605-3p, down-miRs: miR-30c-2-3p, -6828-5p, -22-3p). ST6GAL2 expression was normalized to total protein (Ponceau staining) and divided by normalized NTC for each blot. (**D**) RT-qPCR analysis for samples as in C. All samples are normalized to GAPDH as an endogenous housekeeping gene and then to NTC. All experiments were done in triplicate.

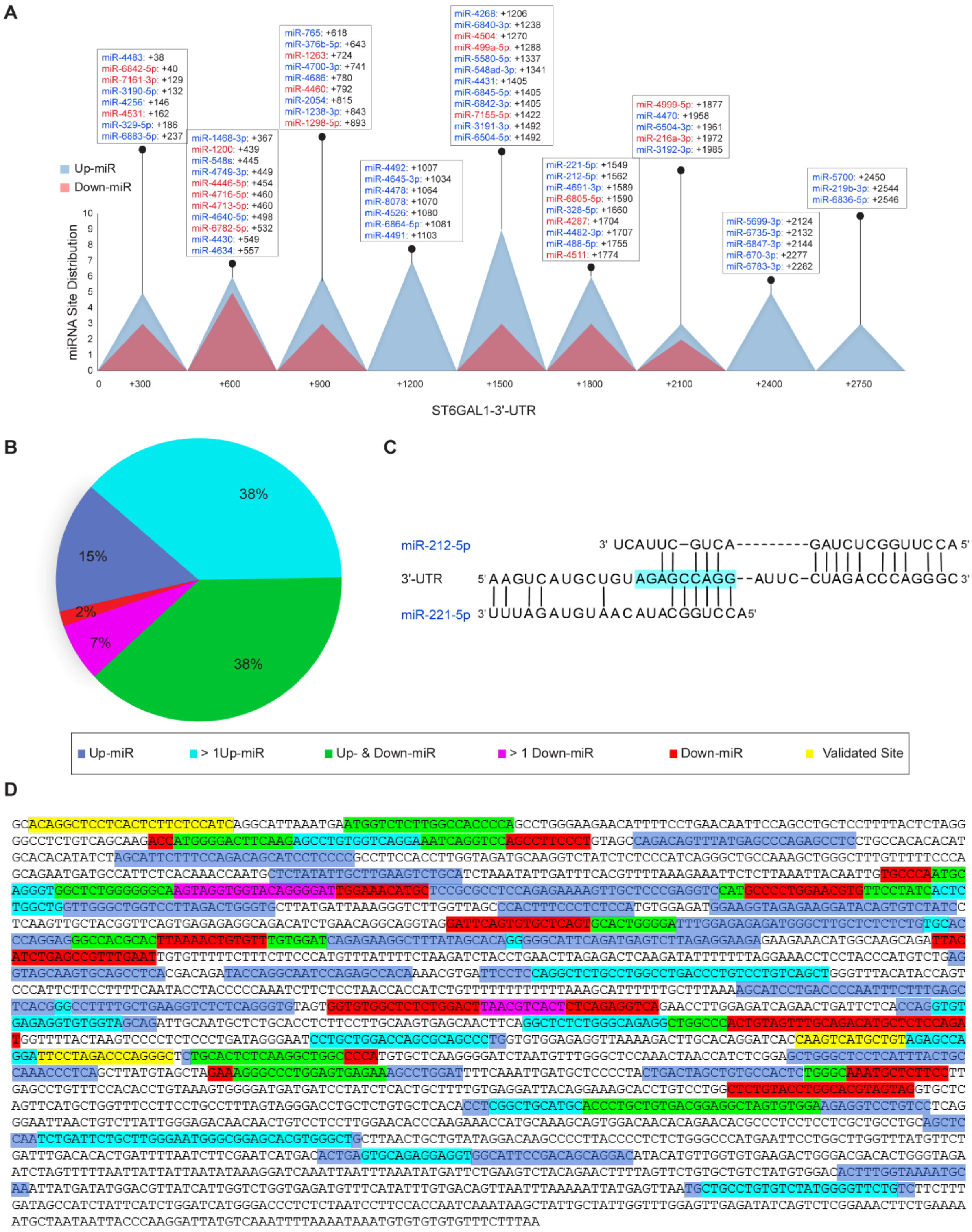
A
miR-4483: +38
miR-6842-5p: +40
miR-7161-3p: +129
miR-3190-5p: +132
miR-4256: +146
miR-4531: +162
miR-329-5p: +186
miR-6883-5p: +237
miR-1468-3p: +367
miR-1200: +439
miR-548s: +445
miR-4749-3p: +449
miR-4446-5p: +454
miR-4716-5p: +460
miR-4713-5p: +460
miR-4640-5p: +498
miR-6782-5p: +532
miR-4430: +549
miR-4634: +557
miR-765: +618
miR-376b-5p: +643
miR-1263: +724
miR-4700-3p: +741
miR-4686: +780
miR-4460: +792
miR-2054: +815
miR-1238-3p: +843
miR-1298-5p: +893
miR-4492: +1007
miR-4645-3p: +1034
miR-4478: +1064
miR-8078: +1070
miR-4526: +1080
miR-6864-5p: +1081
miR-4491: +1103
miR-4268: +1206
miR-6840-3p: +1238
miR-4504: +1270
miR-499a-5p: +1288
miR-5580-5p: +1337
miR-548ad-3p: +1341
miR-4431: +1405
miR-6845-5p: +1405
miR-6842-3p: +1405
miR-7155-5p: +1422
miR-3191-3p: +1492
miR-6504-5p: +1492
miR-221-5p: +1549
miR-212-5p: +1562
miR-4691-3p: +1589
miR-6805-5p: +1590
miR-328-5p: +1660
miR-4287: +1704
miR-4482-3p: +1707
miR-488-5p: +1755
miR-4511: +1774
miR-4999-5p: +1877
miR-4470: +1958
miR-6504-3p: +1961
miR-216a-3p: +1972
miR-3192-3p: +1985
miR-5699-3p: +2124
miR-6735-3p: +2132
miR-6847-3p: +2144
miR-670-3p: +2277
miR-6783-5p: +2282
miR-5700: +2450
miR-219b-3p: +2544
miR-6836-5p: +2546
Up-miR
Down-miR
miRNA Site Distribution
0 +300 +600 +900 +1200 +1500 +1800 +2100 +2400 +2750
ST6GAL1-3'-UTR
B
38%
15%
2%
7%
38%
C
miR-212-5p 3' UCAUUC−GUCA---------GAUCUCGGUUCCA 5'
3'-UTR 5' AAGUCAUGCUGUAGAGCCAGG--AUUC−CUAGACCCAGGGC 3'
miR-221-5p 3' UUUAGAUGUAACAUACGGUCCA 5'
Up-miR
> 1Up-miR
Up- & Down-miR
> 1 Down-miR
Down-miR
Validated Site
D

**Fig. S10.**

**Predicted miRNA binding sites for ST6GAL1 3'-UTR.** (**A**) Map of the binding sites for miRNA identified as hits by our miRFluR assay. Sites shown are the most stable hybridization sites predicted by RNAhybrid (*21*). Annotations are given for every 300 bp. (**B**) Pie chart representing the distribution of miRNA site overlap within the 3'-UTR. Sites are defined as overlapping if the annotated hybridization sites share nucleotides. Periwinkle blue: up-miRs with no overlap, turquoise blue: overlap between 2 or more up-miRs, green: overlap between a set of up-miRs and down-miRs, magenta: overlap between 2 or more down-miRs, red: down-miRs with no overlap. Percent is function of sum of miRNA hits. (**C**) The sequence alignment highlighting the overlap between validated sites of miR-221-5p and -212-5p on 3'-UTR. (**D**) Map of the 3'-UTR with color-coded overlaps among different groups of miRNAs defined in B. Yellow highlights validated sites.

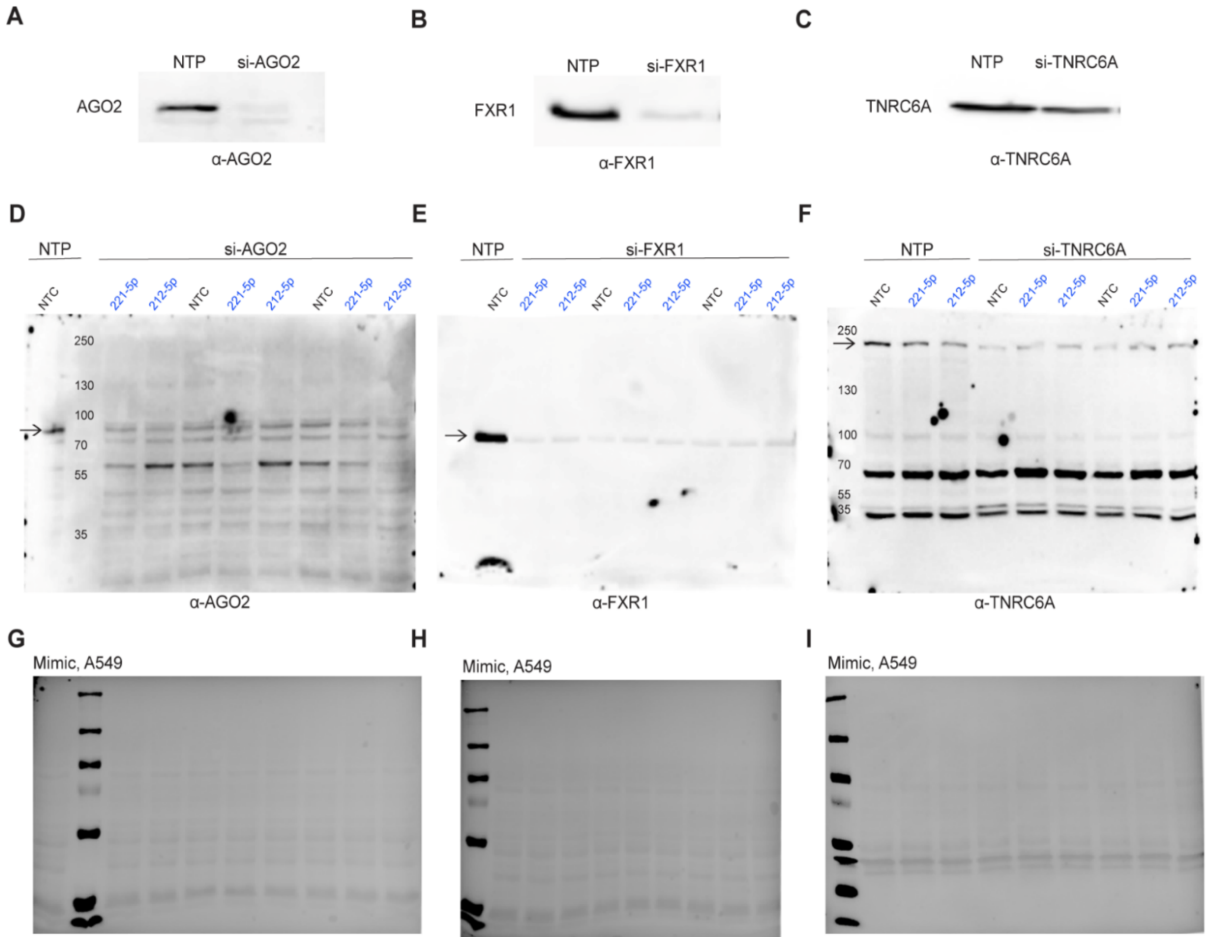


**Fig. S11.**

**Validation of siRNA knock down against AGO2, FXR1 and TNRC6A in A549 human lung carcinoma cell line for experiments shown in Fig. 4.** (**A to C**) Representative Western blots for control experiments verifying knockdown of AGO2 (A), FXR1 (B) and TNRC6A (C), by their respective siRNA pools (50 nM siRNA, 48 h post-transfection). (**D to F**) Western blot analysis confirming knockdown of AGO2 (D), FXR1 (E) and TNRC6A (F) for samples used in Fig. 4 and S12. Arrows indicate the target bands. (**G to I**) Ponceaus of blots shown in D to F, respectively. Experiments shown are biological triplicates.

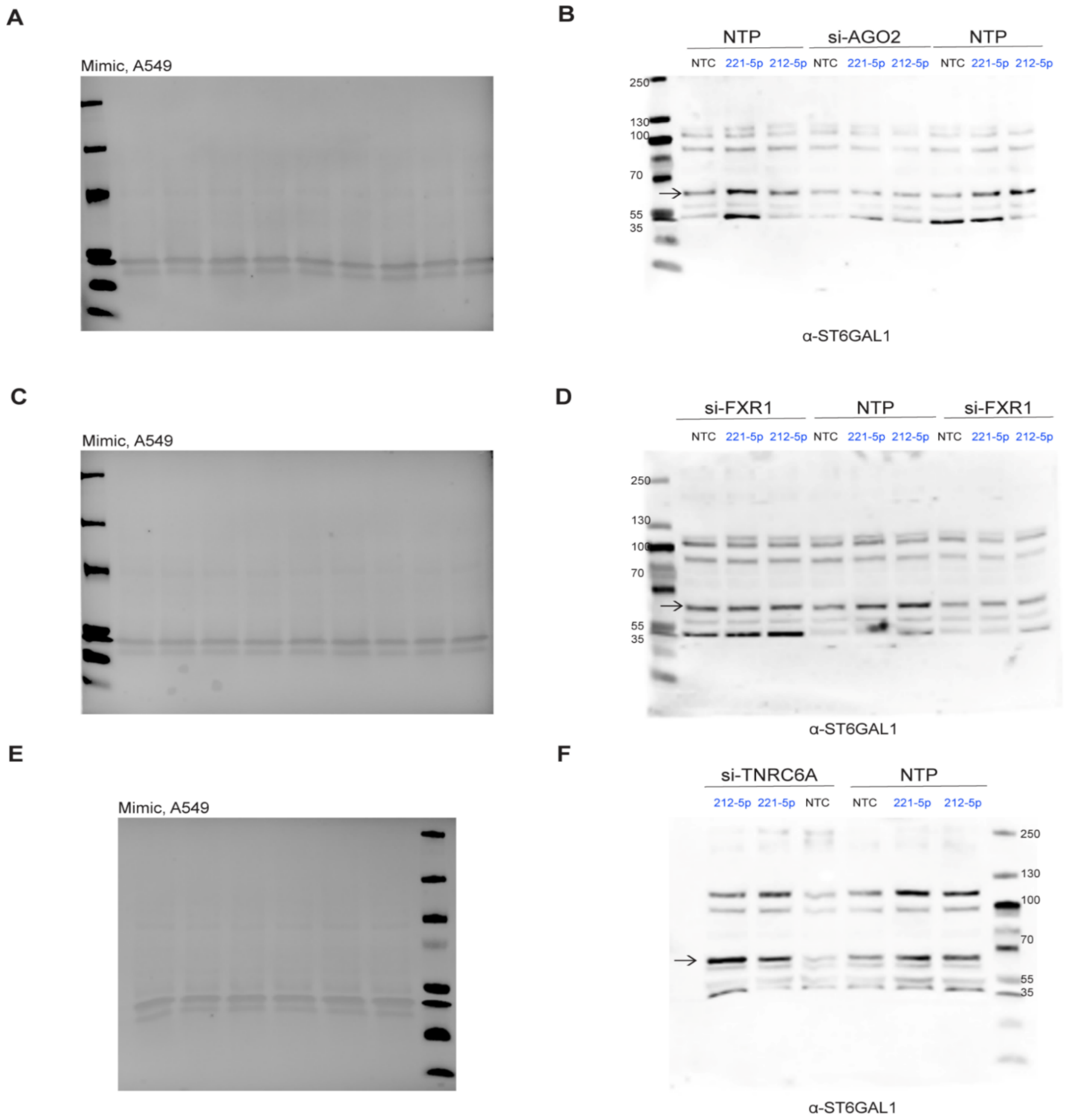


**Fig. S12.**

**Upregulation of expression by miRNAs requires FXR1 and AGO2 in A549 human lung carcinoma cell line.** (**A**) Ponceau of Western blots shown in B. (**B**) Western blot of AGO2 shown in Fig. 4D. (**C**) Ponceau of Western blot shown in D. (**D**) Western blot of FXR1 and NTP shown in Fig. 4D. (**E**) Ponceau of Western blot shown in F. (**F**) Western blot of TNRC6A shown in Fig. 4D. Arrows indicating the validated ST6GAL1 signal.

| Primer Name | Sequence (5' → 3') | Sample |
|---|---|---|
| A. PCR amplification of ST6GAL1 or ST6GAL2 3'-UTR | | |
| ST6GAL1-FWD[i] | CGGACCATTCACTGCTAAG | gDNA, HEK293T |
| ST6GAL1-REV[i] | TTAAAGAAACACACACACATTTATTTTA | gDNA, HEK293T |
| ST6GAL2-FWD | AAAGGGTTTCTTGGGAATC | gDNA, HEK293T |
| ST6GAL2-REV | TTCTAGACAAATGAAAACATG | gDNA, HEK293T |
| B. RT-qPCR[i] quantification of ST6GAL1, ST6GAL2 or GAPDH mRNA | | |
| ST6GAL1-FWD1 | GAACACCCAAGAAACCATGCA | Total RNA, A549 or HT-29 |
| ST6GAL1-REV1 | ACGTGCTCCGCCCATTC | Total RNA, A549 or HT-29 |
| ST6GAL1-FWD2 | AACACCCAAGAAACCATGCAA | Total RNA, PANC1 or OVCAR3 |
| ST6GAL1-REV2 | CGTGCTCCGCCCATTC | Total RNA, PANC1 or OVCAR3 |
| ST6GAL2-FWD | GAAGGAGCCACGTGTTGGA | Total RNA, A549 or HT-29 |
| ST6GAL2-REV | GCGGGTTCAGCATTTTGG | Total RNA, A549 or HT-29 |
| GAPDH-FWD | GGTGTGAACCATGAGAAGTATGA | Total RNA, all cell lines |
| GAPDH-REV | GAGTCCTTCCACGATACCAAAG | Total RNA, all cell lines |
| C. PCR amplification of ST6GAL1 mutant 3'-UTR | | |
| 221-MUTA-FWD | CTCTGCACTCTCAAGGC | 221-MUTA-gBlock |
| 221-MUTA-REV | TTAAAGAAACACACACACATTTAT | 221-MUTA-gBlock |
| 221-MUTB-FWD | CGGACCATTCACTGCTAAG | 221-MUTB-gBlock |
| 221-MUTB-REV | TCTAGGAATGGACCGTCTACT | 221-MUTB-gBlock |
| 212-MUTA-FWD | ATGATTCTGAAGTCTACAGAAC | 212-MUTA-gBlock |
| 212-MUTA-REV | TTAAAGAAACACACACACATTTAT | 212-MUTA-gBlock |
| 212-MUTB-FWD | CGGACCATTCACTGCTAAG | 212-MUTB-gBlock |
| 212-MUTB-REV | AGCTCCGAGATGGTTAGTTTG | 212-MUTB-gBlock |
| 4531-MUT-FWD | CAGGCATTAAATGAATGGTCTCT | pFmiR-ST6GAL1 |
| 4531-MUT-REV | TTAAAGAAACACACACACATTTAT | pFmiR-ST6GAL1 |

[i] FWD, forward; REV, reverse; RT-qPCR, Reverse transcription quantitative polymerase chain reaction.

**Table S1.**
Primer sequences for PCR amplification of W.T. (A) or mutant 3'-UTRs (C) or RT-qPCR quantification of mRNAs (B).

**Data S1. (separate excel file)**
**miRFluR results for ST6GAL1 and ST6GAL2**. Data analysis before and after thresholds.
